# The effect of G-quadruplexes on TDP43 condensation, distribution, and toxicity

**DOI:** 10.1101/2024.04.30.591873

**Authors:** Emily G. Oldani, Kevin M. Reynolds Caicedo, McKenna E. Spaeth Herda, Adam H. Sachs, Erich G Chapman, Sunil Kumar, Daniel A. Linseman, Scott Horowitz

## Abstract

The events that lead to protein misfolding diseases are not fully understood. Many proteins implicated in neurodegenerative diseases (e.g., TDP43) interact with nucleic acids, including RNA G-quadruplexes. In this work, we investigate whether RNA G-quadruplexes play a role in TDP43 condensation in biophysical and cellular models. We find that G-quadruplexes modulate TDP43 aggregation *in vitro* and condensation in multiple cell types, including yeast, HEK293T, and motor-neuron-like NSC-34 cells. In yeast cells, treatment with G-quadruplexes causes increased TDP43 accumulation in cells before cellular death. In HEK293T cells expressing TDP43, incubation with G-quadruplex-binding small molecules causes an increase in G4 stability that also stabilizes TDP43 and reduces TDP43 condensation induced by proteasomal or oxidative stress. Finally, in NSC-34 cells overexpressing exogenous TDP43, we show that G-quadruplexes co-localize with TDP43 condensates under stress conditions and treatment with G-quadruplex-binding small molecules decreases TDP43-mediated toxicity. Together, these findings suggest that a novel future approach to explore for treating protein misfolding diseases may be to target specific RNA structures such as G-quadruplexes.

## INTRODUCTION

Trans-active response DNA binding protein 43 kDa (TDP43), is involved in numerous steps of RNA metabolism and regulation for many transcripts^1–5^. It is primarily present and active in the nucleus but can be found in the cytoplasm, even in healthy cells^6–8^. TDP43 inclusions are found in multiple neurodegenerative and protein misfolding diseases, including cystic fibrosis, amyotrophic lateral sclerosis (ALS), and frontotemporal lobar degeneration (FTLD)^1,2,9–16^. Aggregation of wild type (WT) TDP43 is a hallmark of sporadic amyotrophic lateral sclerosis, which accounts for nearly 90% of ALS cases, yet the mechanism of this aggregation remains unknown^17^.

Based on the RNA-binding capacity of TDP43, using RNA to chaperone TDP43 has been tried in several contexts. For example, TDP43 cytotoxicity was rescued by the expression of human long non-coding RNA (lncRNA) NEAT1 in *Saccharomyces cerevisiae* and in HEK293T cells^17–19^. In yeast, the deletion of debranching enzyme DBR1, and the resulting cytoplasmic build-up of lariat introns, has been observed to reduce the toxicity of TDP43^20^. Additionally, short oligonucleotides with well-characterized TDP43 binding capacity, such as Clip_34nt, have been shown to ameliorate neurotoxic aggregation^21,22^.

In addition to its consensus GU-rich binding sequence, TDP43 has been shown previously to bind G-quadruplexes (G4s)^23^. Quadruplexes are four-stranded helices in which guanine forms tetrads, stabilized by cations located in the center of the tetrad stack. Notably, a SELEX experiment found that the binding to G4s was tightest in cases where the TDP43 could not effectively bind its evolved binding partner, UG-repeat RNA. While the authors suggested that TDP43 could bind and transport G-quadruplex-containing mRNAs into neurites for local translation, it is also possible that the SELEX experiment was to a misfolded form of TDP43 that could then be more representative of disease^24–26^. A study testing the effects of short oligonucleotides on TDP43 aggregation and toxicity suggested G4s could mitigate the toxicity of TDP43 in cells^23^. Recently, it was shown that RNA G4s (rG4s) are formed preferentially in cells under stress conditions, as is likely to occur in neurodegenerative disease^27,28^. In addition to binding TDP43, rG4s have recently been shown to have a strong effect on the aggregation of many proteins^29,30^. Therefore, we sought to explore whether modulating rG4s could affect TDP43 condensation and toxicity.

## RESULTS

### The effect of DNA G4s on in vitro eGFP-TDP43 aggregation

Although it has been previously shown that G4s can bind to TDP43^23^, the ability for G4s to directly modulate TDP43 aggregation was unclear. Therefore, we performed *in vitro* aggregation assays with TDP43 and G4s *(Figure 1a).* To test different states of TDP43 as starting points for aggregation, we isolated both the dimer peak and a >670 kDa peak from the size exclusion chromatography (SEC) at the end of protein purification.

**Figure 1.**
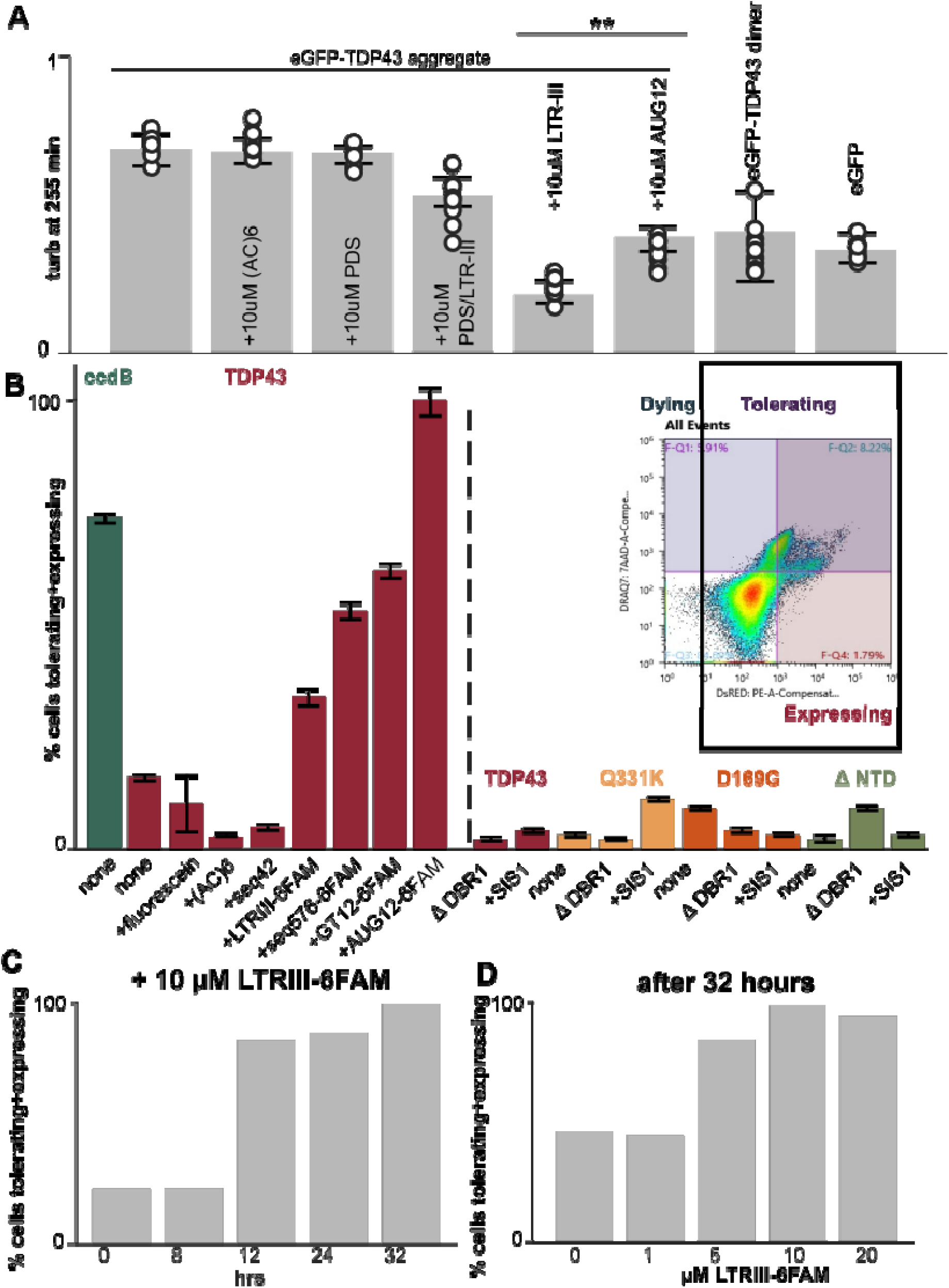
G4s modulate TDP43 aggregation *in vitro* and in yeast. G4s modulate TDP43 aggregation *in vitro* and in yeast. A. The coincubation of eGFP-TDP43 with LTRIII-6FAM slows down the aggregation of the protein as measured by turbidity at 395 nm after 225 minutes. 50 µM eGFP-TDP43 was co-incubated with 10 µM LTR-III and/or 10 µM PDS in 40 mM HEPES pH 7.5, 300 mM NaCl, 1 mM MgCl2, 10% glycerol, 5 mM DTT, for 8.5 hrs at RT and allowed to aggregate. Dimer and >670 kDa peak (“aggregate”) eGFP-TDP43 refer to the sample of protein separated via SEC as being dimeric or approximately 10 times the molecular mass of eGFP-TDP43 monomeric recombinant protein. Each time point has 3 technical and 3 independent replicates. Error bars indicate standard error. One-way ANOVA with Tukey’s comparison test were done where applicable. **p = 0.000001, *p = 0.00703. B-D. The treatment of TDP43-DsRED expressing yeast with nucleic acids results in an increase in the fraction of cells tolerating and expressing the TDP43-DsRED. B. TDP43 and disease point mutants have diminished vitality when expressing TDP43-DsRED, which is rescued by the addition of 10 µM G4 and controls. Overexpression of SIS1 and deletion of DBR1 in yeast do not increase the levels of TDP43 in cells. Inset displays the gating and classification for used for all samples (see Methods for gating rationale) for 10 µM LTRIII-6FAM and TDP43-DsRED expressing yeast after 32 hrs of growth. C. A time course of the normalized % cells tolerating and expressing TDP43-DsRED co-incubated with 10 µM LTRIII-6FAM. D. A dosage curve of the normalized fraction cells tolerating and expressing TDP43-DsRED with LTRIII-6FAM after 32 hrs. 100,000 cells/replicate used for all cytometry. Biological replicate experiments of Figure 1 C, D shown in Supplementary Figure 7.

LTRIII DNA (5’ GGGAGGCGTGGCCTGGGCGGGACTGGGG 3’), a G4 previously demonstrated to have potent chaperone activity^31^, was selected to screen with TDP43 and labeled with 6-Carboxyfluorescein (6FAM) on the 3’ end. The G4 LTRIII-FAM was then mixed with 50 μM eGFP-TDP43 >670 kDa peak sample as separated by SEC (*Supplementary Figure 1*) and monitored for turbidity at 395 nm as an indication of aggregation and disaggregation *(Figure 1a, Supplementary Figure 2*). As a control, eGFP alone was used. When the G4 was added to the 50 μM eGFP, there was little change in the turbidity *(Supplementary Figure 2*). The untreated eGFP-TDP43 aggregate; however, continued to increase in turbidity over time, eventually visibly precipitating out of solution (*Figure 1a, Supplementary Figure 2*). The addition of 10 µM of the G4 LTRIII-6FAM DNA decreased the turbidity of the eGFP-TDP43 aggregate sample and prevented the protein from precipitating out of solution (*Figure 1a, Supplementary Figure 2*). As a further control, we tested this assay with (AC)6, a sequence to which TDP43 does not have affinity that contains no G bases and is not a G4. We did not observe a decrease in the turbidity with (AC)6 (*Figure 1a, Supplementary Figure 2*). Together, these results show that the G4 was able to at least in some cases prevent the aggregation of the recombinant eGFP-TDP43 *in vitro*, and suggested the ability to disaggregate small TDP43 aggregates in solution.

### DNA G4s cause greater TDP43 accumulation in yeast

To test the effects of G4s on TDP43 in cells, we developed a cytotoxicity assay using flow cytometry (*Figure 1b inset*). Overexpression of human WT TDP43 is cytotoxic in yeast^8^. There are known genetic rescues of TDP43 cytotoxicity in yeast; for example, the overexpression of SIS1, and the deletion of debranching enzyme 1 (DBR1)^32,33^. SIS1 is a chaperone protein endogenously expressed in *S. cerevisiae*. DBR1 debranches lariat RNAs; without it, these lariats are not decayed and consequently build up in the cytoplasm^18,33^. These controls decreased TDP43 cytotoxicity in yeast as measured by spot titers (*Supplementary Figure 3*), in a manner that was not dependent on the use of a DsRED tag (*Supplementary Figure 4*)^34^.

For this assay, we used the DNA-binding DRAQ7 far red death stain as a marker for dying cells and the DsRED intensity to quantify the amount of TDP43 in the yeast (see methods). Cells could express high levels of TDP43 and either continue to live, or express high levels of TDP43 and die and show high DRAQ7 staining. We refer to the latter population as “tolerating”, as these cells tolerated a greater level of TDP43 expression before dying (*Figure 1b inset*). This quantification allowed us to monitor the effect of TDP43 levels on the viability of the yeast.

We then used this flow cytometry assay to examine the effects of TDP43 expression level on cytotoxicity in yeast. We quantified the number of cells that were either dying without high TDP43 expression (*Figure 1b inset,* top left quadrant), dying after tolerating high TDP43 expression (*Figure 1b inset,* top right quadrant) or surviving with high TDP43 expression (*Figure 1b inset,* bottom right quadrant). We used this technique to quantify the amount of accumulated TDP43-DsRED, comparing WT protein and two disease point mutants, Q331K and D169G^35^, and domain deletion ΔNTD of TDP43. Of note, when measured by cell cytometer, the previously found controls that rescued toxicity by spotting assay did not increase in the amount of WT TDP43 expressed and tolerated in cells, suggesting that these do not work primarily by increasing tolerated TDP43 levels (*Figure 1b*). As a result, we looked to other potential strategies to stabilize TDP43 in cells using G4s.

Before testing the effects of G4s on TDP43 in yeast, the presence of the 6FAM labeled G4 in the yeast cells was confirmed by confocal microscopy, as was the expression of DsRED-tagged TDP43 (*Supplementary Figure 5*). For comparison with G4s, we tested the effects of two known RNAs that bind TDP43 as part of its regular splicing function, Cy5-AUG12 RNA (5’ GUGUAUUGAUUG 3’) and Cy5-GU12 RNA (5’ GUGUGUGUGUGU 3’), both of which showed an increase in the level of expressed and tolerated TDP43 by flow cytometry.

Disruption of the RRMs by 5 F-> L mutations (F147L and F149L in RRM1, F197L, F22L, and F231L in RRM2), did not result in a cytotoxic yeast model^32^ and had no change from between the cultures treated with the G4 and those without (*Supplementary Figure 6*).

When TDP43-DsRED expressing yeast were co-incubated with G4 DNA displaying previously identified chaperone activity, LTRIII or seq576 (5’ TGTCGGGCGGGGAGGGGGGG 3’)^31^, the amount of TDP43 tolerated and expressed DsRED protein accumulated before cell death increased at similar levels to the evolved TDP43 binders (*Figure 1b*). Treatment of cells with single stranded controls seq42 and (AC)6 did not result in increased TDP43-DsRED tolerance (*Figure 1b*). The amount of TDP43 present in cells also increased as a function of time incubated with the G4 (*Figure 1c)* and as a function of G4 concentration (*Figure 1d*). It may be the case that the increase in TDP43-DsRED being tolerated and expressed in yeast treated with LTRIII (*Figure 1b-d*), is due to increased non-aggregated protein (*Supplementary Figure 5*) in the cytoplasm. This may be a result of TDP43-DsRED forming numerous, large aggregates in yeast in the absence of G4s combined with an overall increase in dying cells (*Figure 1b*, *Supplementary Figure 7*). When there are fewer dying cells, the fraction of cells with TDP43-DsRED expressed and tolerated increases for the cell population (*Figure 1b-d*). Together, these experiments showed that adding G4s could increase the cell’s tolerance for high TDP43 levels. Next, we wanted to test whether targeting G4s intracellularly using small molecules could change TDP43 condensation and prevent toxicity in a more physiological system, and for these experiments we moved to mammalian cells^31,32^

### G-quadruplex-binding small molecules (G4 binders) attenuate the nuclear condensation of TDP43 in HEK293T cell models

After observing that two DNA G4 structures modulate TDP43 aggregation *in vitro* and in yeast model systems, we next examined the effects of G4s on TDP43 condensation in mammalian cell models. We developed an oxidative stress model using HEK293T cells stably overexpressing wild-type (WT) TDP43 tagged with YFP (HEK293T-TDP43-YFP)^40^. Oxidative stress was induced using sodium arsenite (NaAsO_2_), leading to distinct nuclear condensation of the TDP43-YFP (*Figure 2*). This condensation was validated through co-staining with anti-TDP43 antibodies, consistent with previous TDP43 studies^36^ *(Supplementary Figure 8a*). In contrast, no condensates formed in HEK293T cells overexpressing an empty YFP tag under the same conditions (*Supplementary Figure 8b*). Stable overexpression of TDP43-YFP was further validated by Western blot (*Supplementary Figure 8c*). Using live-cell confocal imaging, we observed TDP43 condensation with 4 hrs of NaAsO_2_ treatment (50 μM), with puncta continuing to grow over the 8 hrs experiment (*Supplementary Figure 8d*). These experiments demonstrated a mammalian model for the nuclear condensation of TDP43.

**Figure 2.**
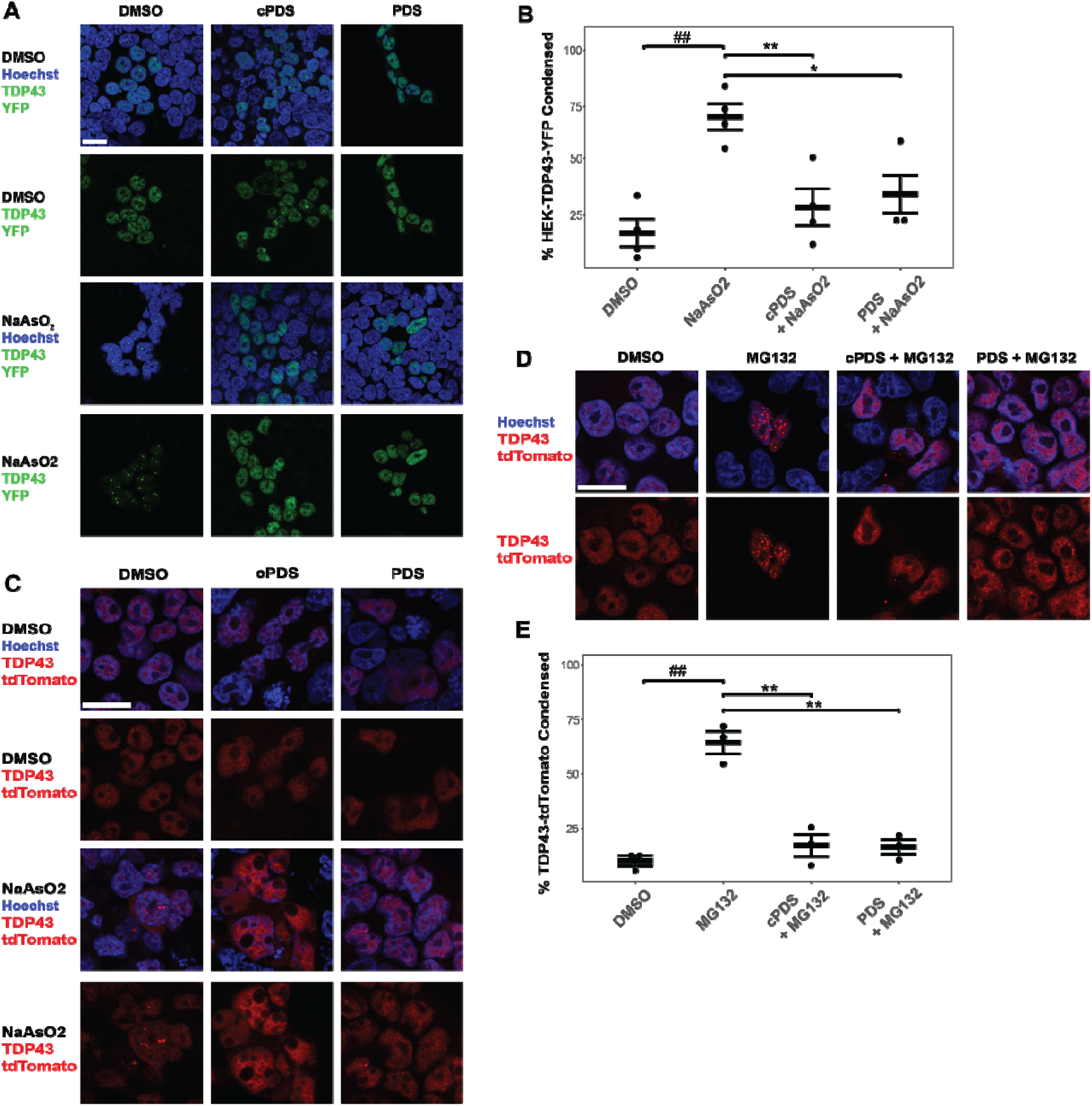
G4 Binders mitigate oxidative and proteasomal stress induced condensation in HEK293T cells overexpressing TDP43. Nuclear stain is Hoechst 33242 (blue) A. Stable overexpression HEK293T-TDP43-YFP (green) cells were treated with 10 µM PDS and cPDS, for 8 hrs prior to oxidative stress (NaAsO_2_, 10 µM, 8 hrs), 40x, scale bar = 20 µm. B. Quantification of condensed nuclear and cytoplasmic TDP43-YFP. Total number of cells containing condensates were divided by total transfected TDP43-YFP expressing cells; (Welch’s two sample t-test, unpaired; ##, **, p < 0.01; *, p <0.05), n = 4. C. Oxidative stress and G4 Binders assays were repeated in HEK293T cells with TDP43-tdTomato (red) transient transfection, 60x, scale bar = 20 µm. D-E. HEK293T cells were subject to proteosome inhibitor MG132 (10 µM) for 8 hrs. Pretreatment with G4 binders (10 µM for 8 hrs) and proteosome stress were repeated and condensates were quantified; (Welch’s two sample t-test, unpaired; ##, **, p < 0.01) n = 3.

Pyridostatin (PDS), a small molecule that binds DNA and RNA G4s, and carboxypyridostatin (cPDS), an analog of PDS that specifically binds rG4s, were employed to test the effects of targeting G4s on TDP43 condensation in cells^27,37–39^. *In vitro*, these molecules on their own did not affect TDP43 condensation (*Figure 1a*), suggesting that observed effects on TDP43 condensation in cells would be due to effects on G4s and/or rG4s in cells. When PDS and cPDS were incubated in WT HEK293T cells, immunocytochemistry (ICC) shows marked increases in the levels of G4s present in cells (*Supplementary Figure 9*). HEK293T-TDP43-YFP cells were pretreated with 10 μM cPDS or 10 μM PDS for 8 hrs prior to the introduction of 10 µM NaAsO_2_. In the presence of either molecule, TDP43 condensation was significantly reduced; instead of puncta formation, a diffuse TDP43-YFP signal was primarily observed within the nucleus (*Figure 2a, b*). To ensure this was not a product of the stable over-expression tag, we transiently transfected TDP43-tdTomato into HEK293T cells, and similar results were seen with PDS and cPDS treatments (*Figure 2c*). As controls, PDS was tested for its ability to directly modulate TDP43 aggregation or whether it blocked the ability for a G4 to affect TDP43 condensation (*Figure 1a, Supplementary Figure 2*), and it was found that it was not sufficient to do either. Together, these results show that adding rG4 binders dramatically reduces TDP43 condensation in HEK293T cells induced under conditions of oxidative stress.

Although NaAsO_2_ stress is commonly used to test intracellular condensation^36,40^, we wanted to confirm whether the effects of the rG4 binding molecules were consistent with another type of protein stress. As a second stress model, we utilized proteasomal inhibition by the addition of MG132^41,42^. As expected, MG132 treatment induced robust nuclear condensation of TDP43. However, cells treated with PDS or cPDS exhibited a diffuse nuclear TDP43 distribution, similar to the unstressed controls (*Figure 2d*). We also treated these cells with the G4 and rG4 antibody BG4^43,44^ to observe rG4 localization. Notably, we observed some cytoplasmic condensation of TDP43 that overlapped with BG4 stain, suggesting the colocalization of RNA G4s with TDP43 (*Supplementary Figure 10*), prompting further evaluation.

### rG4 binders significantly inhibit the cytosolic redistribution and condensation of overexpressed TDP43 in differentiated motor neuron-like NSC34 cells

To evaluate the efficacy of these ligands in a more physiologically relevant system, we expanded these experiments to motor neuron-like NSC-34 cells. Under control conditions, differentiated NSC-34 cells overexpressing WT TDP43-tdTomato consistently displayed nuclear localization of TDP43 (*Figure 3a, left panel, DMSO)*. However, upon exposure to oxidative or proteasomal stress, TDP43 exited the nucleus and formed cytoplasmic condensates (*Figure 3a, middle & right panel; & 3b, middle & right panel columns MG132, NaAsO_2_)*. Notably, TDP43 puncta did not colocalize with stress granule marker TIA-1 (*Supplementary Figure 11a*). Similar to results observed in the HEK293T cell model, preincubation of NSC-34 cells with either cPDS or PDS significantly inhibited the nuclear export of TDP43 induced by either NaAsO_2_ or MG132, resulting in a diffuse TDP43 signal remaining largely within the nucleus (*Figure 3c*).

**Figure 3.**
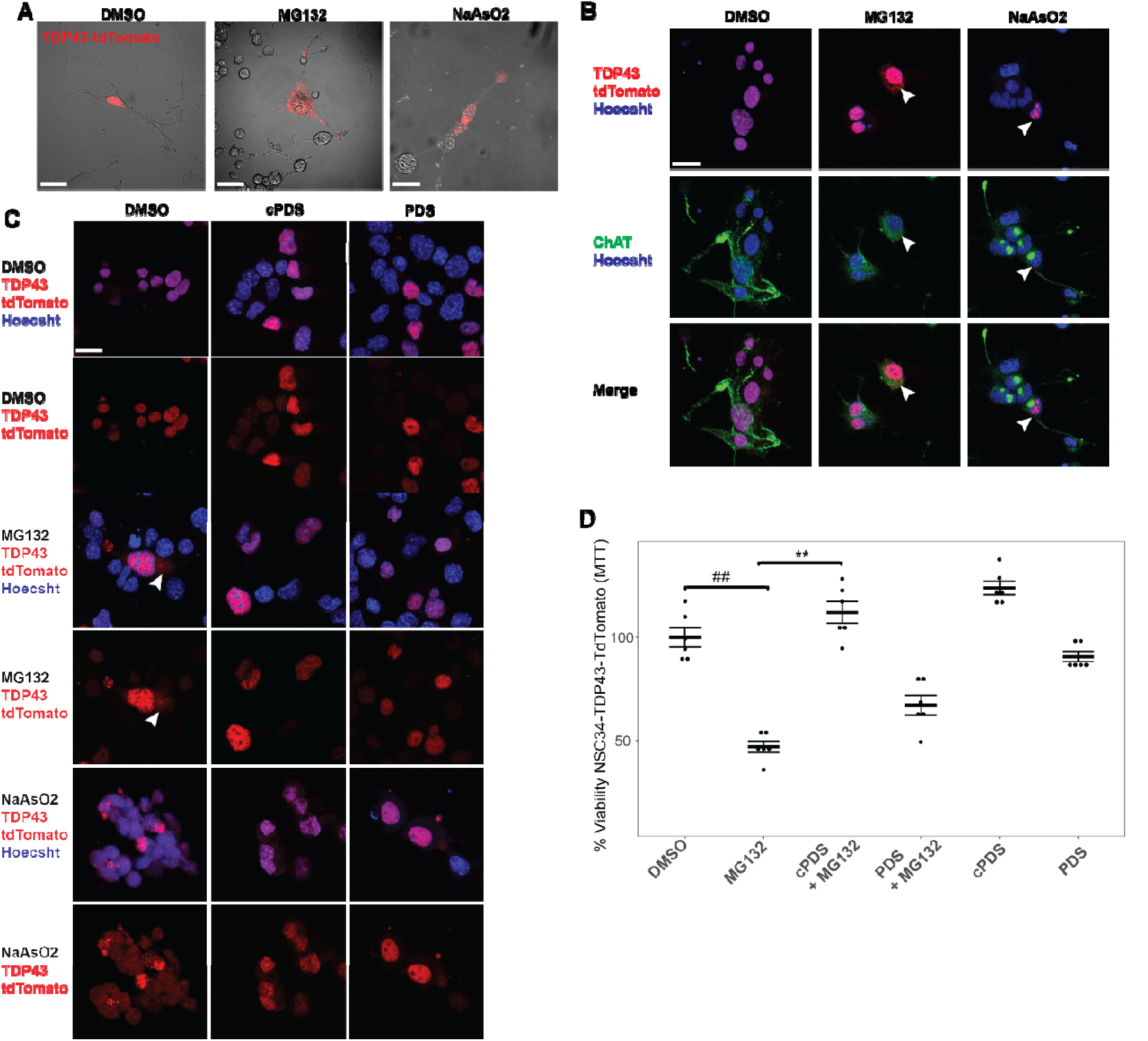
G4 Binders mitigate oxidative and proteasomal stress induced condensation in TDP43 overexpressed NSC34 cells. A. NSC-34 cells transfected with TDP43-tdTomato (red) differentiated for 96 hrs in low serum media with 10 µM Retinoic Acid (R.A.). MG132 and NaAsO_2_ induced cytosolic condensation of TDP43 within 8 hrs. B. Differentiation was confirmed with ICC, 4% PFA, anti-ChAT, ms Alexa-Fluor 647 (green), nuclear labeled Hoechst 33342 (blue), and cytosolic inclusions are formed within 8 hrs of 5 µM NaAsO_2_ and 5 µM MG132 stress. C. When treated with G4 binders, cytosolic condensates are lessened in both stress conditions; n = 6, scale bar = 20 µM. D. MTT data of NSC34-TDP43-tdTomato; in overexpression model, cell viability and survival significantly improved when pretreated with 5 µM cPDS and near significance in 5 µM PDS (Two-way ANOVA, Tukey’s post-hoc test, unpaired; ##, **, p < 0.01) n = 6.

To assess the impact of the rG4 binders on cell death when TDP43 is overexpressed in the NSC-34 cells, cell viability was quantified using a standard MTT assay. TDP43-tdTomato overexpression was shown to have no appreciable effect on cell viability under non-stressed conditions when compared to controls overexpressing an empty vector (tdTomato plasmid with no TDP43) or WT NSC-34 cells (*Supplementary Figure 11b*). Without TDP43 overexpression, cells showed minimal loss of viability and no puncta formation, regardless of the presence or absence of proteasomal stress or rG4 binders (*Supplementary Figure 11c, d*). However, cell death increased substantially when TDP43 was overexpressed, particularly under proteasomal stress. Treatment with the rG4 binder cPDS significantly preserved cell viability, while PDS showed a smaller, non-significant effect, suggesting that rG4s may play a more prominent role than G4s in stabilizing TDP43 (*Figure 3d*).

### G-quadruplexes localize to TDP43 condensates under induced cellular stress

Mechanistically, the ability for G4 binders to affect TDP43 condensation is most plausible if both TDP43 and G4s co-localize in the described cell models. To test whether this was the case, we used an immunostaining assay for RNA G4s using a BG4 antibody in HEK and NSC-34 cells. Cells were fixed with either paraformaldehyde (PFA) or methanol to appreciate any differences in localization of G4s, as PFA fixation causes rRNA staining in the cytoplasm while methanol fixation allows both rG4 and G4 staining in the cytoplasm and nucleus^45^. As previously shown, addition of cPDS/PDS increases the level of G-quadruplexes in cells (*Supplementary Figure 9*). rG4s co-localize with TDP43 condensates both intra- and perinuclearly after oxidative stress or proteasome inhibition (*Figure 4c, d)*. HEK-TDP43-YFP cells fixed with PFA observed similar overlap when perturbed with oxidative stress (data not shown). Both fixation stainings revealed that G4/rG4 levels increased as a function of both the NaAsO_2_ and proteasomal stress (*Figure 4c, d*), consistent with previous observation of increased rG4 formation under several stress conditions^27^. This co-localization is consistent with the direct modulation of TDP43 aggregation by G4s as observed *in vitro*, and the effects of the G4 binders observed here.

**Figure 4.**
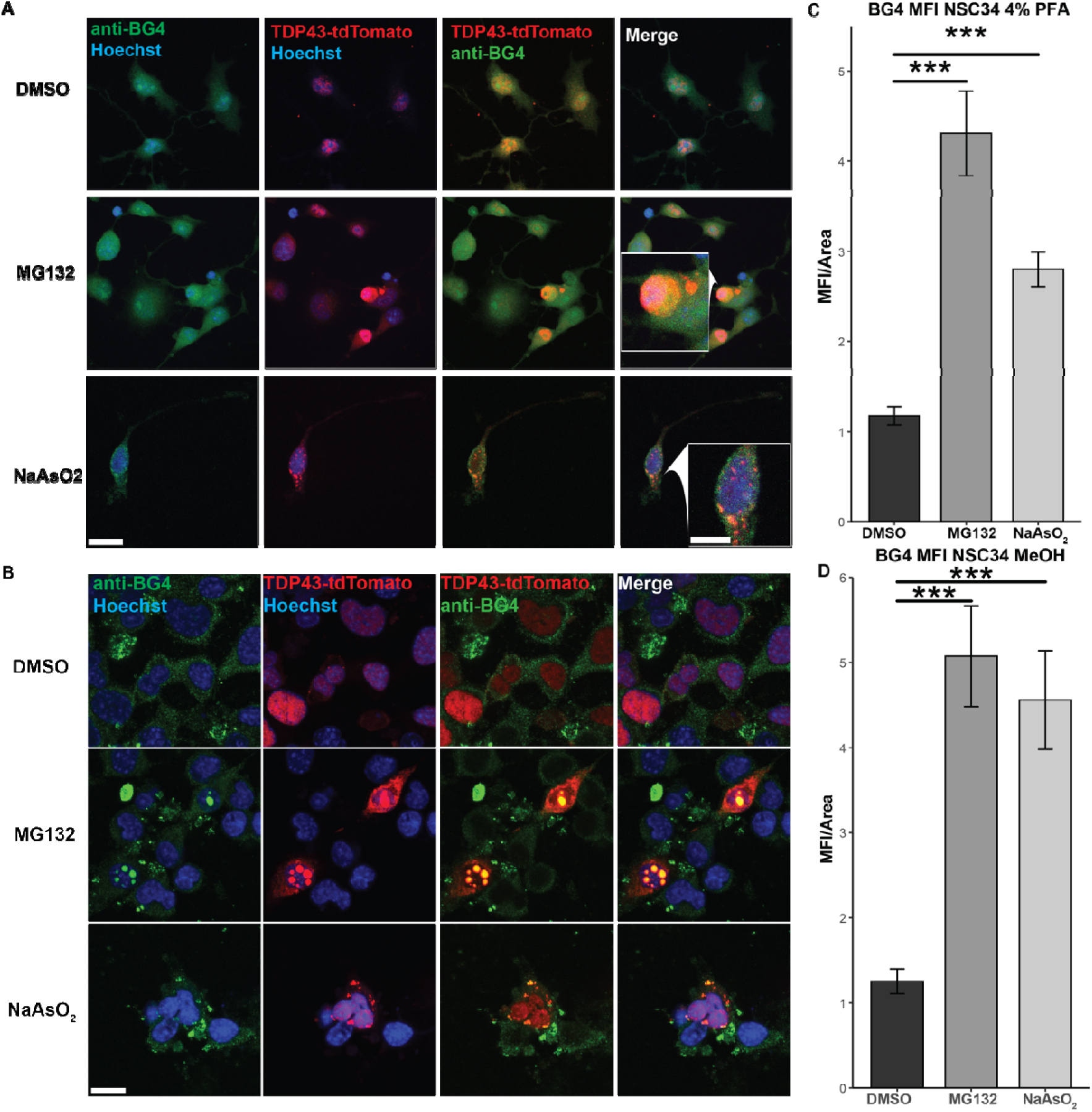
A, B. Immunocytochemistry NSC34 TDP43-tdTomato cells stained with anti-BG4 (1:50) suggested colocalization of RNA G-Quadruplexes with condensed TDP43 under oxidative stress conditions. Immunocytochemistry performed with two fixations, ice-cold 4% PFA (A) and ice-cold 100% methanol (B). Scale bars, anti-BG4/Hoechst = 20 µM; insert = 10 µM. C, D. Quantifications of mean fluorescent intensity of both fixation methods, respectively (C and D), Overall intensities are less in PFA fixation than MeOH, likely due to nuclear inclusions that are largely excluded in the PFA fixation. (Two-way ANOVA, Tukey’s post-hoc test, unpaired; ***, p < 0.001; n = 3)

## DISCUSSION

In this study, we examined the ability of rG4s to modulate the condensation and toxicity of TDP43. *In vitro* experiments showed that G4s prevented the continued aggregation of pre-formed TDP43 aggregates. Experiments in yeast showed that exogenously added G4s could affect the level of TDP43 accumulation before cell death. Experiments in HEK293T cells demonstrated that the addition of rG4-binding molecules could prevent the formation of TDP43 condensates. Finally, experiments in motor neuron-like NSC-34 cells showed that rG4s colocalized to TDP43 condensates, and that the addition rG4 binding molecules both ameliorated condensation and rescued viability from both oxidative and proteasomal stress. Together, these results suggest that modulating rG4s merits further investigation as therapeutic approach for diseases involving TDP43 mislocalization and aggregation.

Markedly, we do not see TDP43 colocalize with stress granule marker, TIA-1. Our system uses arsenite and MG132 in NSC34 cell models at 2.5 μM and 5 μM, respectively, to elucidate a slower stress response to FL WT TDP43. In systems with higher doses of stress (0.5 mM arsenite for 30 minutes), WT FL TDP43 is shown to create smaller SGs than certain TDP43 mutants^46,48^or TDP43 C-terminal truncation^50,52,53–55^. We speculate that the overexpressed exogenous WT FL TDP43 is not localizing to the SGs in the cytosol due to this lower concentration of stress, as previously shown by *Colombrita et al*^54^.

cPDS is likely to bind rG4s of differing structures with differing affinities. Therefore, our analysis here is a simplification of what is a complex mixture of rG4 structures that are differently affected by the binding molecule. While in some cases, these molecules stabilize rG4 formation^27,37–39^, this effect is likely structure-dependent. It is notable that cPDS was more effective at rescuing cytotoxicity than PDS, suggesting that the primary modulation here was of rG4s and not G4s.

Similarly, there are several different possibilities for the mechanism by which PDS and cPDS affect TDP43. It may be the case that they affect the structural pool of rG4s. As rG4s of parallel and anti-parallel topology have vastly different effects on aggregation^29,47^, the PDS and cPDS could be shifting the population of the rG4 structures in a favorable manner. Similarly, it is possible that the PDS and cPDS are preventing certain deleterious rG4s from binding to TDP43 by competitive binding to the rG4s, although our *in vitro* experiments suggest this possibility is less likely. Because the population of differing structural rG4s topologies in the cell is, to our knowledge, completely unknown, it is difficult to determine which of these factors are most dominant in cells.

Overall, the results here demonstrate proof-of-principle results that modulating rG4 populations is a powerful way to modulate TDP43 aggregation and toxicity. As TDP43 is found to aggregate in multiple diseases^1–3,6–12,15,16^, this could be a promising avenue for further therapeutic study, especially as rG4s are specifically targetable with small molecules. Of note, TDP43 is not the only disease-related protein that binds to G4s, and other neurodegenerative disease aggregation-prone proteins bind G4s as well^30^, suggesting that this strategy could be a beneficial next step in exploring modulating the aggregation of proteins in many diseases.

## STAR Methods

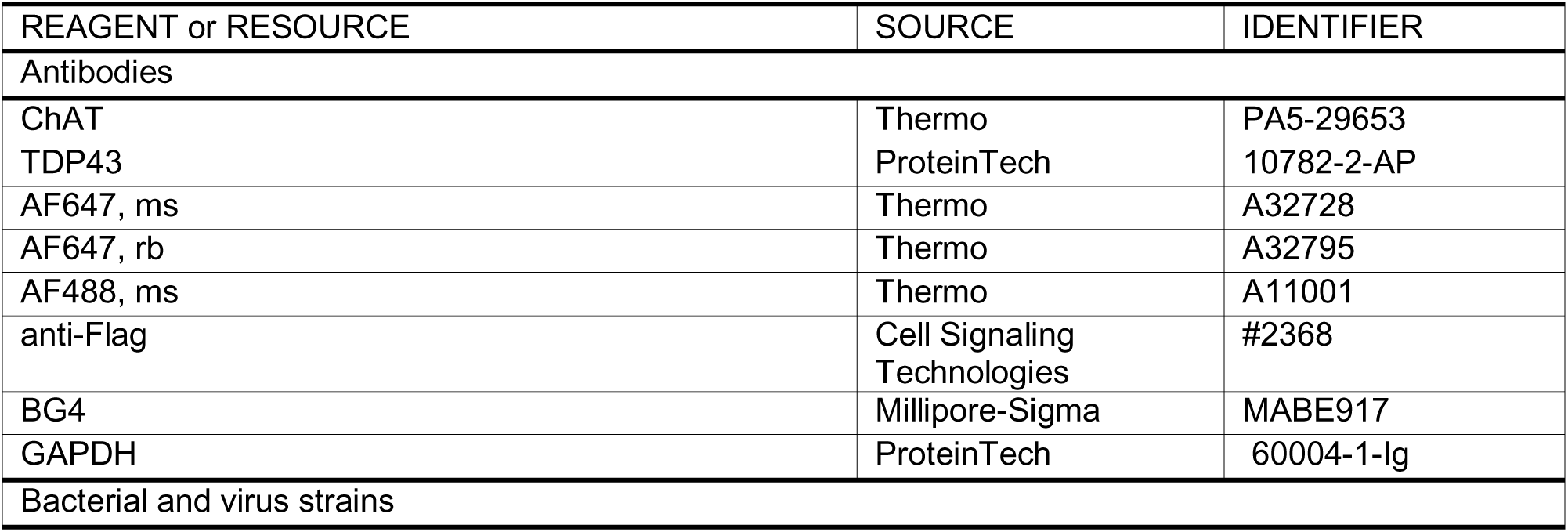
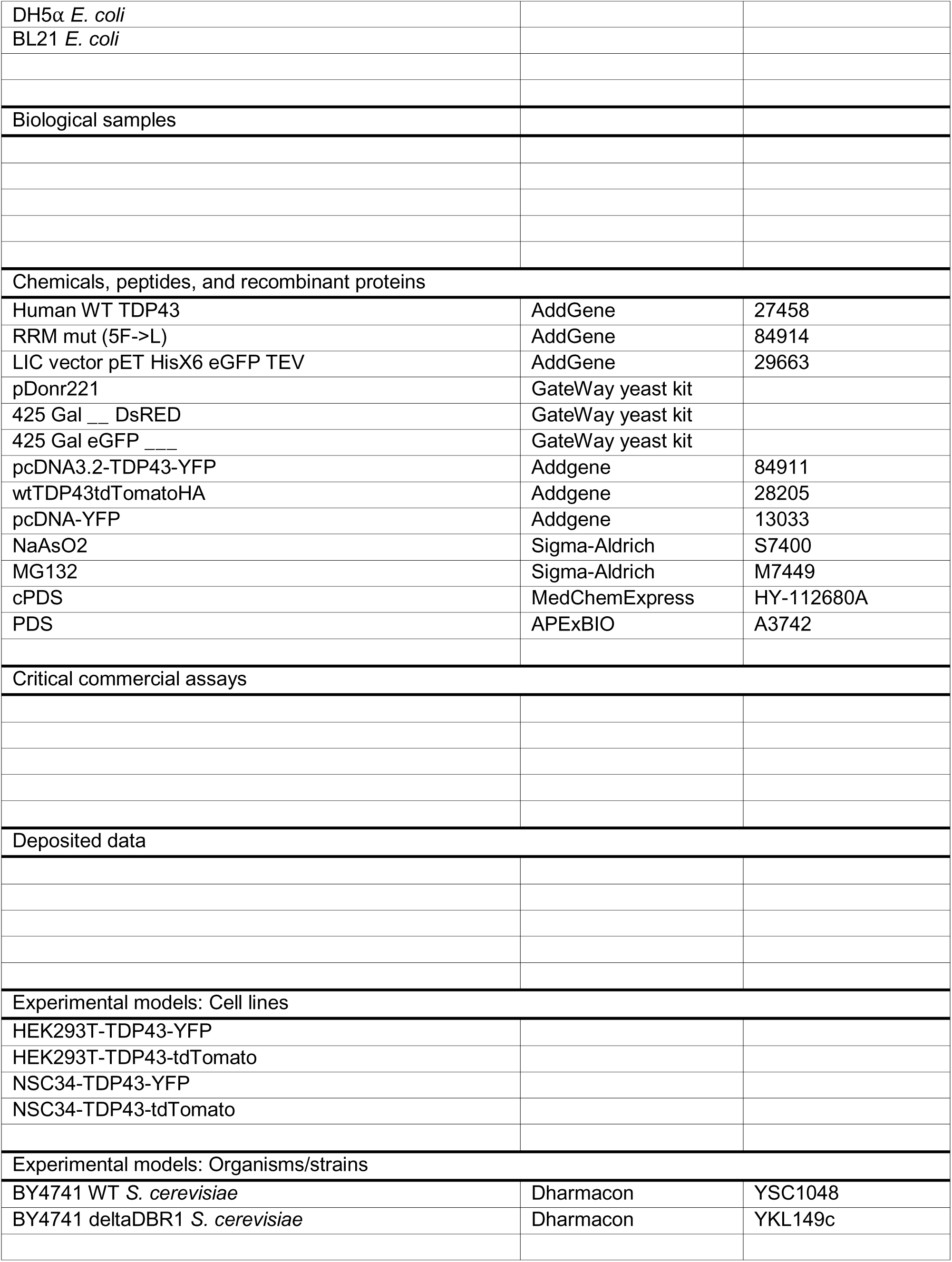
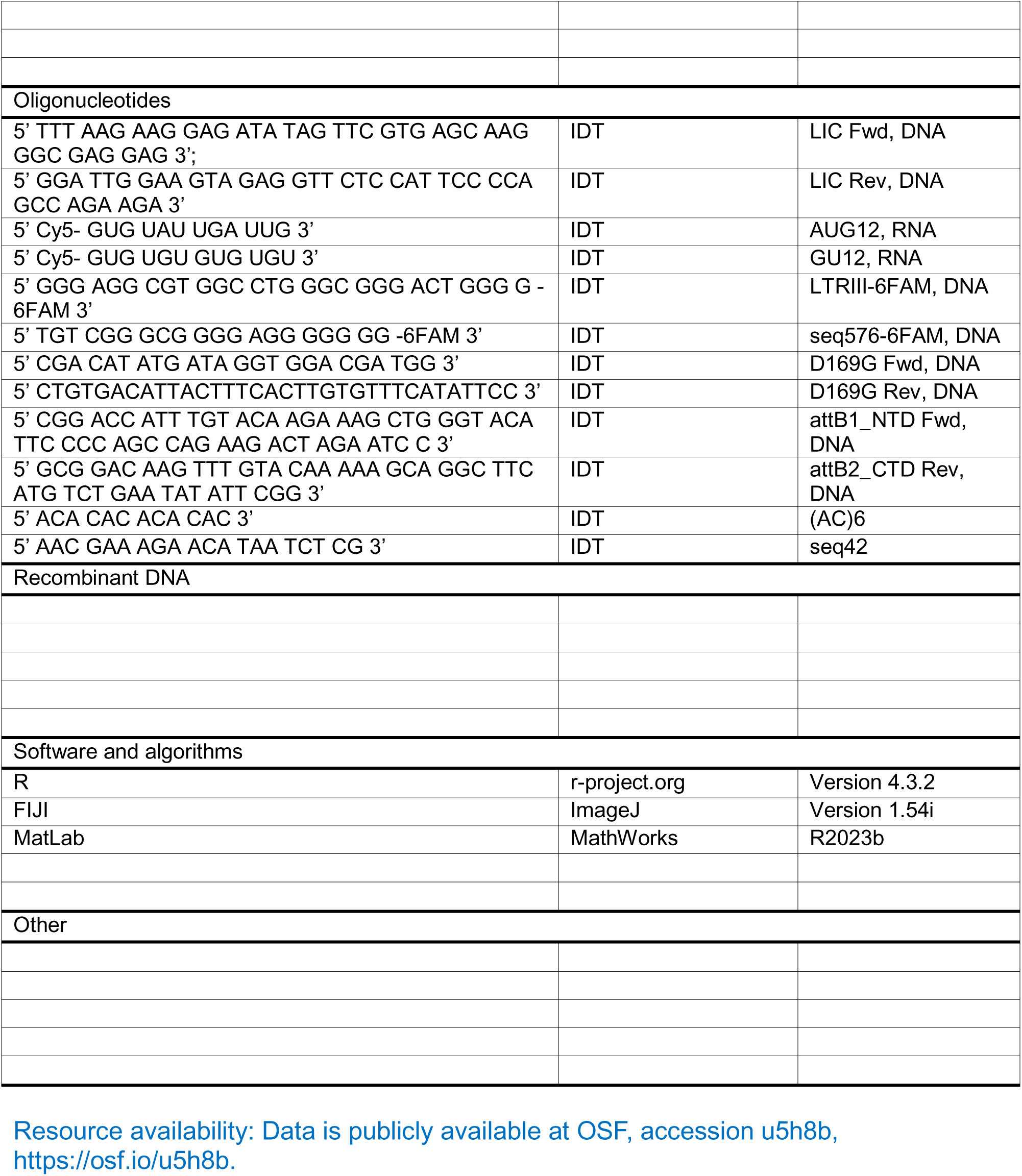

### Cloning

A variety of cloning techniques for recombinant protein expression by *E. coli* and *S. cerevisiae* were used to purify protein for the use of *in vitro* assays and to express hTDP43 in a model organism for *in vivo* assays.

### Ligation Independent Cloning (LIC)

Human WT TDP43 (AddGene #27458), and RRM mut (5F → L) (AddGene #84914) open reading frames (ORFs) were cloned into the LIC vector pET HisX6 eGFP TEV cloning vector from AddGene (AddGene #29663) using QB Berkley LIC protocol with primers designed using the tool from QB3 MacroLab (Fwd: 5’ TTTAAGAAGGAGATATAGTTCGTGAGCAAGGGCGAGGAG 3’; Rv: 5’ GGATTGGAAGTAGAGGTTCTCCATTCCCCAGCCAGAAGA 3’). The primers were used to amplify the ORFs via polymerase chain reaction (PCR), and the LIC vector was digested with SspI restriction enzyme. The synthesized plasmid was then transformed into DH5α *Escherichia coli* before being isolated from 10 mL, single colony cultures, using a Genesee Miniprep kit, and sent for sequencing at Quintara Bio, and ultimately transformed into BL21 expression *E. coli*.

### Site-Directed Mutagenesis (SDM, QuickChange)

The point mutation D169G was generated using the New England Biolabs Q5® Site Directed Mutagenesis Kit (Fwd: 5’ CGA CAT ATG ATA GGT GGA CGA TGG 3’; Rv: 5’ CTG TGA CAT TAC TTT CAC TTG TGT TTC ATA TTC C 3’). Synthesized plasmid was then transformed into DH5α *E. coli* for sequencing at Quintara Bio, and ultimately transformed into BL21 *E. coli* for expression and subsequent purification.

### Gateway Cloning

The TDP43 sequences used were lifted using PCR with primers that included attB sites for compatibility with pDonr221 (Fwd: 5’ CGG ACC ATT TGT ACA AGA AAG CTG GGT ACA TTC CCC AGC CAG AAG ACT AGA ATC C 3’; Rv 5’ GCG GAC AAG TTT GTA CAA AAA GCA GGC TTC ATG TCT GAA TAT ATT CGG 3’). Yeast expression vectors were created using the Gateway® Cloning system and its included protocols^49,51,53^. The plasmid selected was a high copy, leucine auxotrophy, galactose inducible, C-terminal DsRED tagged one, after first being cloned, from a gBlock or PCR product, into a Gateway® Cloning entry vector, pDonr221.

### Yeast cloning, plasmid

BY4741 WT yeast obtained from Dharmacon were grown to an approximate OD_600_ of 0.35, 250 μL were pelleted, rinsed with MBG H2O, briefly suspended in 100 mM LiOAc, before being pelleted and resuspended in 50 μL MBG H2O. 1 μg desired plasmid was then added to the yeast and allowed to incubate for 5 minutes before the addition of 10 μL sheared salmon DNA. 500 μL PLATE (40% PEG-3350, 100 mM LiOAc, 10 mM Tris pH 7.5, 1 mM EDTA) was then added. Samples were vortexed and allowed to incubate at 42°C for 35 minutes, then plated on selection media.

### Protein purifications

The proteins used in these experiments were purified from recombinant DNA expressed by *E. coli* using affinity columns. Buffers, conditions, and purification and growth processes are detailed below for each of the experimentally used proteins.

### eGFP-TDP43 recombinant expression and purification of native protein

Cultures grown from a single colony of BL21 bacteria containing the plasmid for the desired construct were incubated at 37°C to an OD_600_ of approximately 0.4 before being induced with 1 mM IPTG for 6 hrs at 20°C. Cultures were then pelleted at 4°C at 8000 rpm for 30 minutes before decanting the media and resuspending the pellets in approximately 60 mL lysis buffer (40 mM HEPES pH 7.5, 300 mM NaCl, 1 mM MgCl_2_, 50 mM Imidazole, 5 mM βMe, 10% glycerol), DNase (20 μg/μL), CHAPS (1%), and Egg White Lysozyme (0.25 mg/mL) were added and the resulting suspension was stirred at 4°C for 1 hr. After pelleting at 4°C at 31000 RCF for 75 minutes, the supernatant was filtered using a 0.22 μM Genesee bottle top filter before being loaded onto an 8 mL Bio-rad nickel immobilized metal affinity column (IMAC) column in an NGC System from Bio-rad. The column was equilibrated in the lysis buffer and after loading the lysate by sample pump, 5 column volumes of lysis buffer were run over the column, followed by a gradient to 100% elution buffer (40 mM HEPES pH 7.5, 300 mM NaCl, 1 mM MgCl_2_, 500 mM Imidazole, 5 mM βMe, 10% glycerol) over 3 column volumes. Samples were collected from the fractions that corresponded to a peak at 280 nm which occurred between 50-70% elution buffer. Collected fractions were pooled, filtered, and run on a GE Life Sciences Superdex 200 16/600 SEC with SEC buffer (40 mM HEPES pH 7.5, 300 mM NaCl, 5 mM DTT, 10% Glycerol). Purified fractions were pooled and then either used immediately or flash frozen and stored at -80°C for later use.

### Analysis by Size exclusion chromatography

Samples were run on a GE Life Sciences Superdex 200 16/600 SEC with SEC buffer (40 mM HEPES pH 7.5, 300 mM NaCl, 5 mM DTT, 10% Glycerol). Commercial sizing standards (https://www.bio-rad.com/en-us/sku/1511901-gel-filtration-standard?ID=1511901) were run as well.

### Turbidity

The protein was loaded into a 96 well plate and the turbidity of the sample at 395 nm and 600 nm was monitored over time at room temperature with no shaking. 4 measurements were taken of each well, and averaged, every 2 minutes for 90-1200 minutes.

### Yeast viability assays

TDP43 and control protein expressing yeast were examined for yeast vitality by serial dilution on an agar plate. The working concentration of the oligos as drugs for cell sorter experiments was determined to be 10 µM for the low copy, galactose inducible TDP43-DsRED yeast expression plasmid.

### Spotting assay

BY4741 WT yeast containing the Gateway plasmid 425 Gal-ccdB/TDP43-DsRED with the various point mutants and other listed proteins were allowed to grow in 2% glucose selection media (-Leu) to an OD_600_ between 0.1 and 0.3. ODs were measured in triplicate and averaged before being diluted to 0.01 and 7 serial dilutions of 1 to 8 were done. This was spotted onto 2% glucose -Leu agar plates and 4% galactose -Leu agar plates. Plates were imaged every 12 hrs of growth at 30°C for 3 days. The high-copy plasmid, 425, was used.

### Cell sorter analysis

Cell sorting of BY4741 WT 415 Gal TDP43 DsRED live cell imaging. Low copy plasmid is 1 copy (CEN)^53,55^ high copy is 40-60 copies (2µ)^49,53^. AUG12 and GT12 are known TDP43 binders^22^. Cells were grown to an OD_600_ of 0.3 in SD 2% glucose media, then induced with 4% galactose –His media for 8 hrs, then 1 μM LTRIII-3’ 6FAM (5’ GGG AGG CGT GGC CTG GGC GGG ACT GGG G-6FAM 3’), seq576-6FAM (5’ TGT CGG GCG GGG AGG GGG GG 3’), AUG12-3’ 6FAM (5’ GUG UGA AUG AAU-6FAM 3’), GT12-6FAM (5’ GTG TGT GTG TGT-6FAM 3’), or fluorescein was added and cells were allowed to grow for 16 hrs. Cells are pelleted and washed twice with phosphate buffered saline (PBS: 137 mM NaCl, 2.7 mM KCl, 4.3 mM Na_2_HPO_4_,1.47 mM KH_2_PO_4_, pH 7.4), and then resuspended in PBS, stained with DRAQ7 as a nuclear death stain. Excess DRAQ7 is then removed by washing cells twice with PBS before being resuspended in PBS. An SH800 Sony Cell Sorter is then calibrated using a mix of 3 μm and 10 μm nanobeads (Sony Automatic Setup Beads LE-B3001), before sorting 100,000 cells for the following distributions: DsRED (PE-A) vs. Back scattering, DRAQ7 (7AAD) vs. back scattering, 6FAM (CSFE) vs. back scattering, 6FAM vs. DRAQ7, DsRED vs. DRAQ7.

Quadrants for dying, tolerating, and expressing TDP43-DsRED were identified and assigned by creating gated quadrants to separate the high and low levels of DRAQ7 and DsRED. The low levels of DRAQ7 were determined to be anything below a secondary, small population that was above the other cells in the sample along the DRAQ7 axis, as the cells in the low region were the same in the presence and absence of DRAQ7. The low and high levels of DsRED expression were determined by bisecting the population of cells along the DsRED axis. Dying cells were considered to be those in the quadrant of high DRAQ7 and low DsRED. Tolerating cells were considered to be those in the quadrant of high DRAQ7 and high DsRED. Expressing cells were considered to be those in the quadrant of low DRAQ7 and high DsRED. Cells with both low DRAQ7 and low DsRED were not included in the total number of cells.

### Spinning disc confocal microscopy of yeast

BY4741 WT yeast containing the Gateway plasmid 425 Gal-ccdB/TDP43-DsRED with the various point mutants and other listed proteins. The yeast were grown to an OD_600_ of 0.1 in 2% Glucose selection media (-Leu) before being pelleted, resuspended in the same volume 4% Galactose selection media, and allowed to continue to grow for 7.5 hrs. The yeast were then pelleted, washed with PBS twice, resuspended in PBS, and plated for imaging with an Olympus FV3000 spinning disc confocal. Images are processed by using FIJI and are de-speckled and smoothed using the included software of FIJI. Z stacks were averaged and combined for the DsRED channel using the program included in FIJI. G-quadruplex LTRIII -6FAM was observed to be taken into the cytoplasm of the yeast when added directly to the media, as was fluoresceine (*Supplementary Figure 6*).

### Cell culture

Human embryonic kidney 293T (HEK293T) cells were grown in culture medium (High Glucose DMEM (Hy-Clone), 10% HI FBS (Gibco), 1% L-glutamine (Gibco), 1% Pen/Strep (Gibco)) on 10 cm tissue culture dishes and incubated at 37°C and 5% CO_2_. For all experiments, cells between passages 5 and 20 were used to maintain consistency.

Immortalized neuroblastoma x spinal cord cells (NSC-34; Cedarlane Labs, Burlington, ON. CA) were cultured in medium containing high glucose DMEM, 10% FBS, and 1% pen/strep, in 6-well tissue culture plates (CellTreat) and incubated at 37°C and 5% CO_2_.

### Mammalian TDP43 stable and transient overexpression transfection

pcDNA3.2 TDP43 YFP (Addgene #84911) was graciously provided by the Aaron Gitler Lab, wtTDP43tdTOMATOHA (Addgene #28205) was graciously provided by the Zuoshang Xu Lab, and pcDNA3-YFP (Addgene #13033) was graciously provided by the Doug Golenbock Lab. These plasmids were transformed into DH5a *E. coli* before being isolated from 5 mL, single colony cultures, using an Omega Bio-Tek Miniprep Kit. These plasmids were transfected into HEK293T and NSC34 cell lines with Lipofectamine 2000 (Invitrogen, 11668027) 24 hrs prior to stable selection or transient experimentation, respectively.

HEK293T cells were seeded in a 24-well tissue culture plate (CellTreat) at 1.5 x10^5^ and incubated overnight. Lipofectamine 2000 was used per manufacturer’s protocol with 500 ng of pcDNA3.2 TDP43 YFP^40^ added per well. Cells were split the following day into single colonies and cultured in 500 µg/mL G418 to select for YFP positive cells. TDP43-YFP cells were passed at 80% confluency, seeded at 1 x 10^6^ HEK293T cells per passage, and maintained in 200 µg/mL G418. wtTDP43-tdTomatoHA^56^ was transiently transfected as described above.

### Western Blotting

10 µg of HEK293T WT, YFP and TDP43-YFP overexpression cell lysates were separated on a 10% Bolt Bis-Tris Mini Gel (Thermo Fisher, California, United States) and transferred on to PVDF membrane (Thermo Fisher, California, United States) using an iBlot 3 Transfer System (Thermo Fisher, California, United States). Blot was blocked in 1% BSA blocking buffer, incubated in primary antibody for 1 hrs at RT with rabbit anti-TDP43 (ProteinTech, 10782-2-AP, 1:1000), then Donkey anti-rabbit IgG (H+L) Highly Cross-Adsorbed Alexa-Fluor Plus 647 (Invitrogen, A32795, 1:1000) for 1 hrs at RT. GAPDH (ProteinTech, 60004-1-Ig, 1:50000) and Goat anti-mouse IgG (H+L) Highly Cross-Absorbed Alexa-Fluor Plus 488 (Invitrogen, A11001, 1:1000). Bound antibodies were visualized on Bio-rad ChemiDoc Imaging system at 647 nm.

### NSC34 Cell Culture Differentiation

For immunocytochemistry assays, NSC-34 cells were seeded on 0.5 mg/mL Poly-D-Lysine treated #1.5 cover glass in 24 well plates (MatTek) at 75,000 cells/mL. For viability (MTT) assays, NSC-34 cells were seeded at 150,000 cells/mL in 6 well plates (CellTreat). Maintenance media (DMEM, 10% FBS, 1% P/S) was swapped 24 hrs after seeding to differentiation media containing: 1:1 DMEM/F-12 (Ham), 0.5% FBS, 1% modified Eagle’s medium nonessential amino acids (NEAA), 1% P/S, and 10 µM all-trans retinoic acid (atRA), and differentiation media was switched every 48 hrs for 5 days. These cells were cultured and transfected with the wtTDP43tdTOMATOHA plasmid and differentiated with culture media containing 10 μM retinoic acid for 96 hrs before experiments were performed. Optimal differentiation was verified by immunocytochemistry for ChAT (Invitrogen, PA5-29653) antibody staining under normal, oxidative, and proteasomal stress conditions.

### Mammalian Cell TDP43 Condensation

To induce TDP43 condensation, Sodium Arsenite (NaAsO_2_, Sigma-Aldrich, S7400) and MG132 (Sigma-Aldrich, M7449) were added to G4 binder pretreated and negative control prepared HEK293T and differentiated NSC34 cells.

For immunocytochemistry, in HEK293T models, NaAsO_2_ and MG132 were added at 10 µM working concentrations and incubated at 37°C for 8 hrs prior to fixation. In NSC34 models, NaAsO_2_ and MG132 were added at 2.5 µM and 5 µM, respectively, and incubated at 37°C for 8 hrs prior to fixation.

For cell viability (MTT) assays, condensation agents were incubated at 37°C for 36 hrs and concentrations stated above. Addition of NaAsO_2_ to WT HEK293T cells increased endogenous TDP43 mean fluorescence (*Supplementary Figure 12*).

### Administration of G4 Binders

To test neuroprotective effects of G4 binders, Pyridostatin and Carboxypyridostatin were incubated prior to perturbations for 8 hrs at 37°C, then fixed and imaged. Cell models were prepared as previously stated. HEK293T models were treated with 10μM of PDS and cPDS, while NSC34 models were treated with 5 µM. 0.1% and 0.05% DMSO were used as vehicle controls for G4 binder delivery. Addition of cPDS to WT HEK293T cells increased TDP43 mean fluorescence (*Supplementary Figure 12*).

### MTT Tetrazolium Cell Viability Assay

The MTT (3-(4,5-dimethylthiazol-2-yl)-2,5-diphenltetrazolium bromide) tetrazolium reduction assay was used to quantify viability of NSC34 cell lines. Thiazolyl Blue Tetrazolium Bromide (MTT Powder) (Sigma-Aldrich, M2128) reagent was prepared dissolving MTT in Dulbecco’s Phosphate Buffer Solution (DPBS), pH = 7.4, to 5 mg/mL, filtered. 200 µL of reagent was added to differentiated NSC34 plates to a final concentration of 0.45 mg/mL and incubated at 37°C for 3 hrs. 2 mL of DMSO was added to dissolve formazan crystals and agitated on plate shaker for 10 minutes to ensure complete solubilization. Wells were aliquoted into black wall 96 well plate and absorbance were recorded at 570 nm on a Tecan Infinite M200Pro.

### Immunocytochemistry and Live Cell Confocal Imaging

Cells and test compounds were prepared as previously stated and were fixed in ice cold 4% Paraformaldehyde (PFA) for 20 minutes or ice cold 100% Methanol (MeOH) for 10 minutes, permeabilized with solution of 0.1% Triton X-100 and 1x DPBS for 10 min, then blocked with blocking buffer solution (0.2% gelatin, 0.5% Triton X-100, DPBS, pH=7.4) for 1 hr. Primary and secondary antibodies were diluted in blocking buffer and incubated on coverslips away from light at 25°C for 1 hr, respectively (anti-TDP43 (1:50), Rb, ProteinTech, 10782-2-AP; anti-ChAT (1:200), Rb, Invitrogen, PA5-29653; anti-BG4 (1:50), Ms, Millipore-Sigma, MABE917; anti-Flag (MA4) (1:50), Ms, Cell Signaling Tech, #2368; Goat anti-mouse IgG (H+L) Highly Cross-Adsorbed Alexa-Fluor Plus 647 (1:1000), Invitrogen, A32728; Donkey anti-rabbit IgG (H+L) Highly Cross-Adsorbed Alexa-Fluor Plus 647 (1:1000), Invitrogen, A32795). Nuclei were labeled by the addition of 1 ug/mL Hoechst stain (33342, Invitrogen, #62249), incubated away from light for 10 minutes, rinsed with PBS and stored for imaging. Coverslips were imaged via confocal microscopy (Olympus FV3000). For live cell imaging, MatTek 24-well plates were removed for incubator and imaged for a maximum of 30 minutes before being returned to the incubator. Mean fluorescence intensity was calculated by circling cell bodies and/or populations with the “Freehand selections” tool in FIJI and normalized by dividing the mean intensity by the area of the cells.

### Statistical Analysis

Experiments were performed using duplicate or triplicate replicates for all biologic and *in vitro* samples. Error bars represent the mean +/- the standard error. Statistical analyses were performed using Welch’s two sample t-test or two-way analysis of variance (ANOVA) and Tukey’s Post-Hoc test in R and MatLab. Differences between treatment groups were statistically significant at a p value of p < 0.05.

## Acknowledgements

This work was funded by CDMRP ALSRP grants W81XWH-22-1-0218 and HT94252410122. The authors would like to thank David Robinson and Nicole Toro for aid in some of the molecular cloning, and Yan Qin and Anna Dischler for assistance with time lapse microscopy.

## Declaration of Interests

No conflicts of interest

---

**Supplementary Figure 1.**
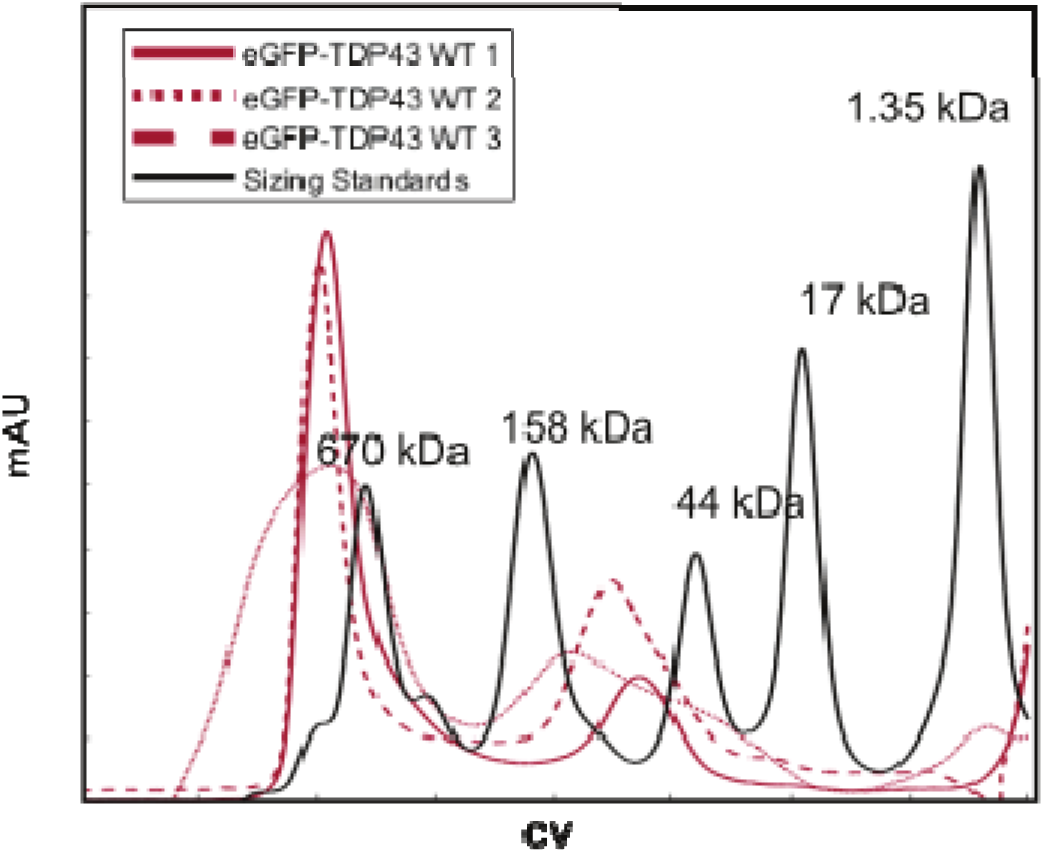
Size exclusion chromatography was used to separate the aggregate eGFP-TDP43 and the dimeric eGFP-TDP43 in 40 mM HEPES pH = 7.5, 300 mM NaCl, 1 mM MgCl_2_, 10% glycerol, in triplicate with Bio-rad sizing standards.

**Supplementary Figure 2.**
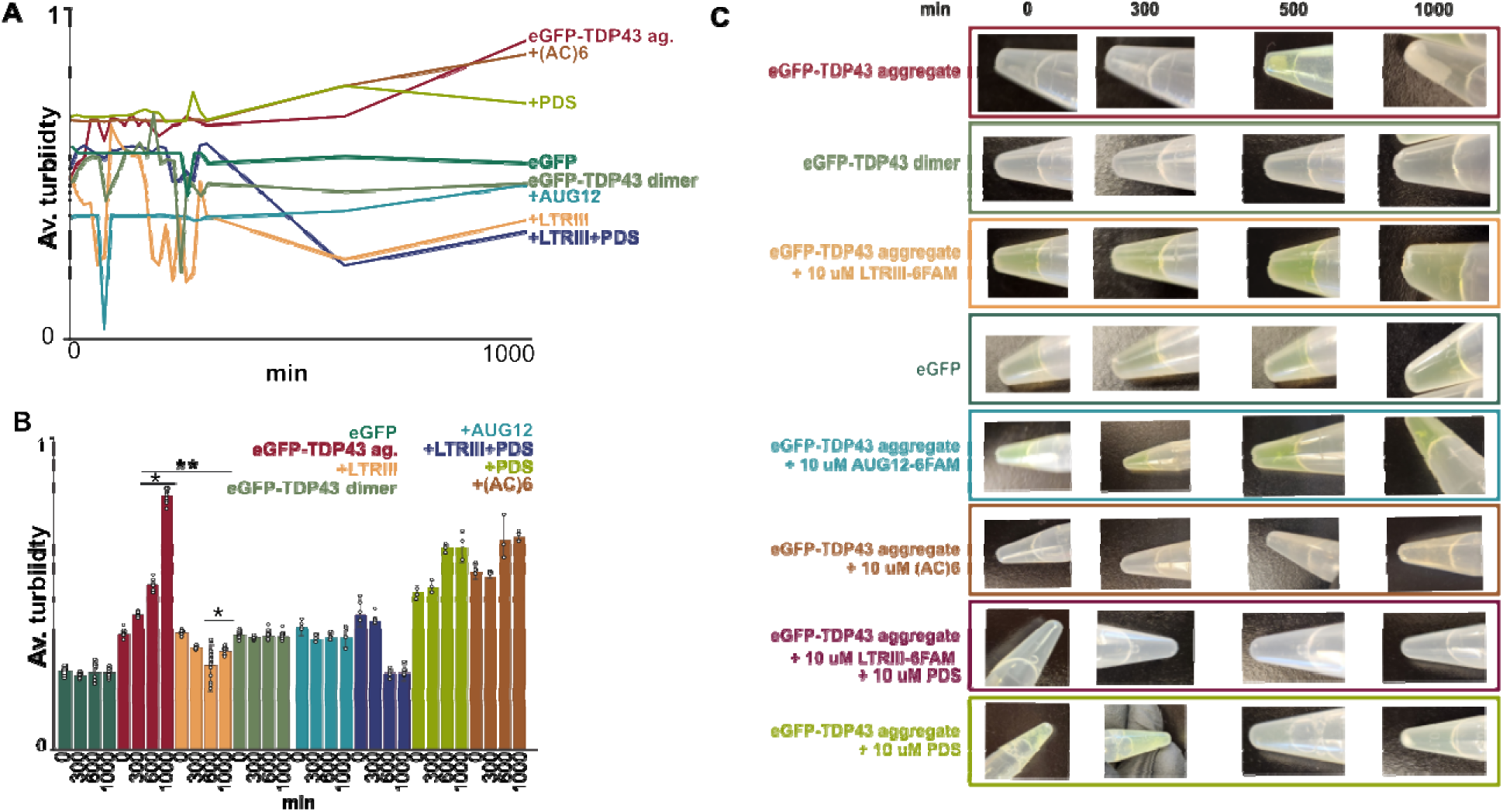
As eGFP-TDP43 aggregate sample (approximately 10X the monomer molecular mass as measured by SEC, <670 kDa) continues to aggregate over time, it begins to precipitate out of solution. A, B. The turbidity over time of the eGFP-TDP43 samples as measured by OD at 395nm by a plate reader over time while shaking on a thermomixer at 37°C. All are aggregate eGFP-TDP43 unless otherwise indicated. C. Photos of the samples at times 0, 300, 500, and 1000 minutes are included to show eGFP-TDP43 aggregate falling out of solution.

**Supplementary Figure 3.**
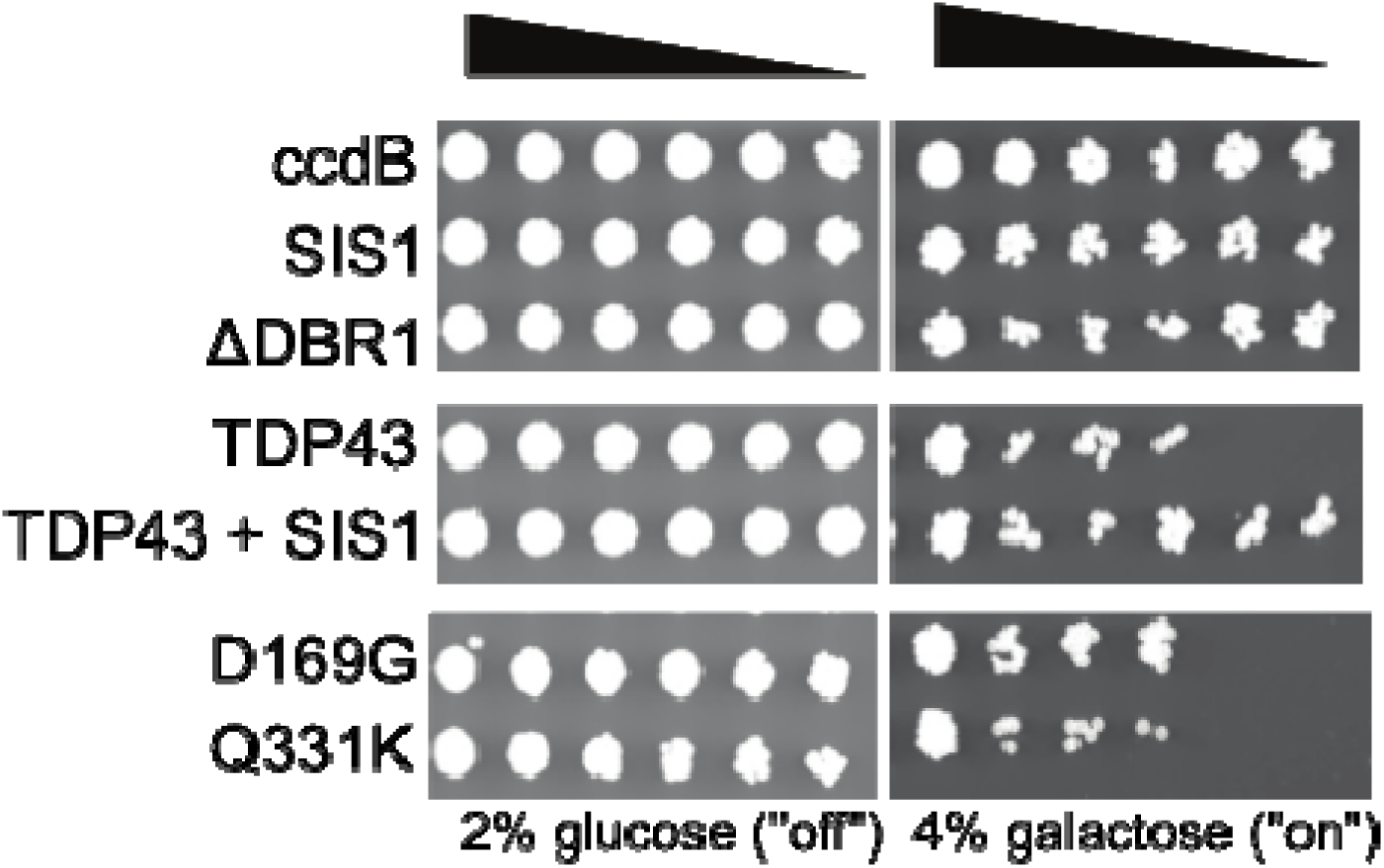
Spotting assay demonstrates the recovery of growth in the cytotoxic TDP43-DsRED expressing yeasts by the over expression of SIS1, and the deletion of DBR1. TDP43 WT and point mutants are expressed in BY4741 WT and ΔDBR1 yeast purchased from Dharmacon using a galactose inducible plasmid with a C-terminal DsRED tag. Serial dilutions of yeast cultures grow in non-toxic protein constructs when expressed. SIS1 is co-expressed using a constitutively active N-terminal GFP plasmid.

**Supplemental Figure 4.**
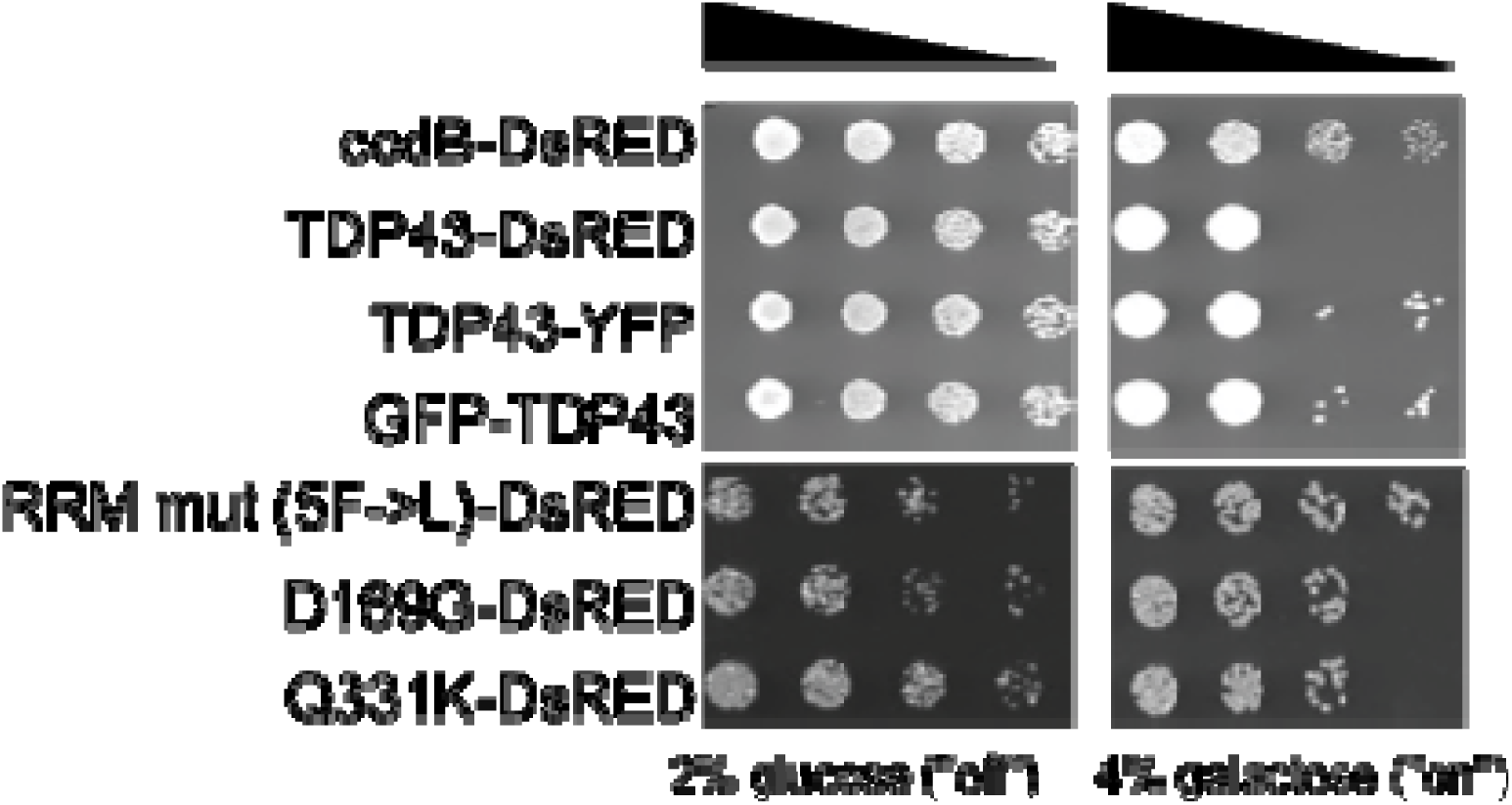
The fluorescent protein fused to TDP43 can be on the N- or C-terminal end, and more than one type of fluorescent protein can be used and still retain a cytotoxic phenotype in yeast. TDP43 WT and point mutants are expressed in BY4741 WT and ΔDBR1 yeast purchased from Dharmacon using a galactose inducible plasmid with a C-terminal DsRED tag. Serial dilutions of yeast cultures grow in non-toxic protein constructs when expressed. The location and identity of the fluorescent protein did not make the TDP43 non-toxic in yeast.

**Supplementary Figure 5.**
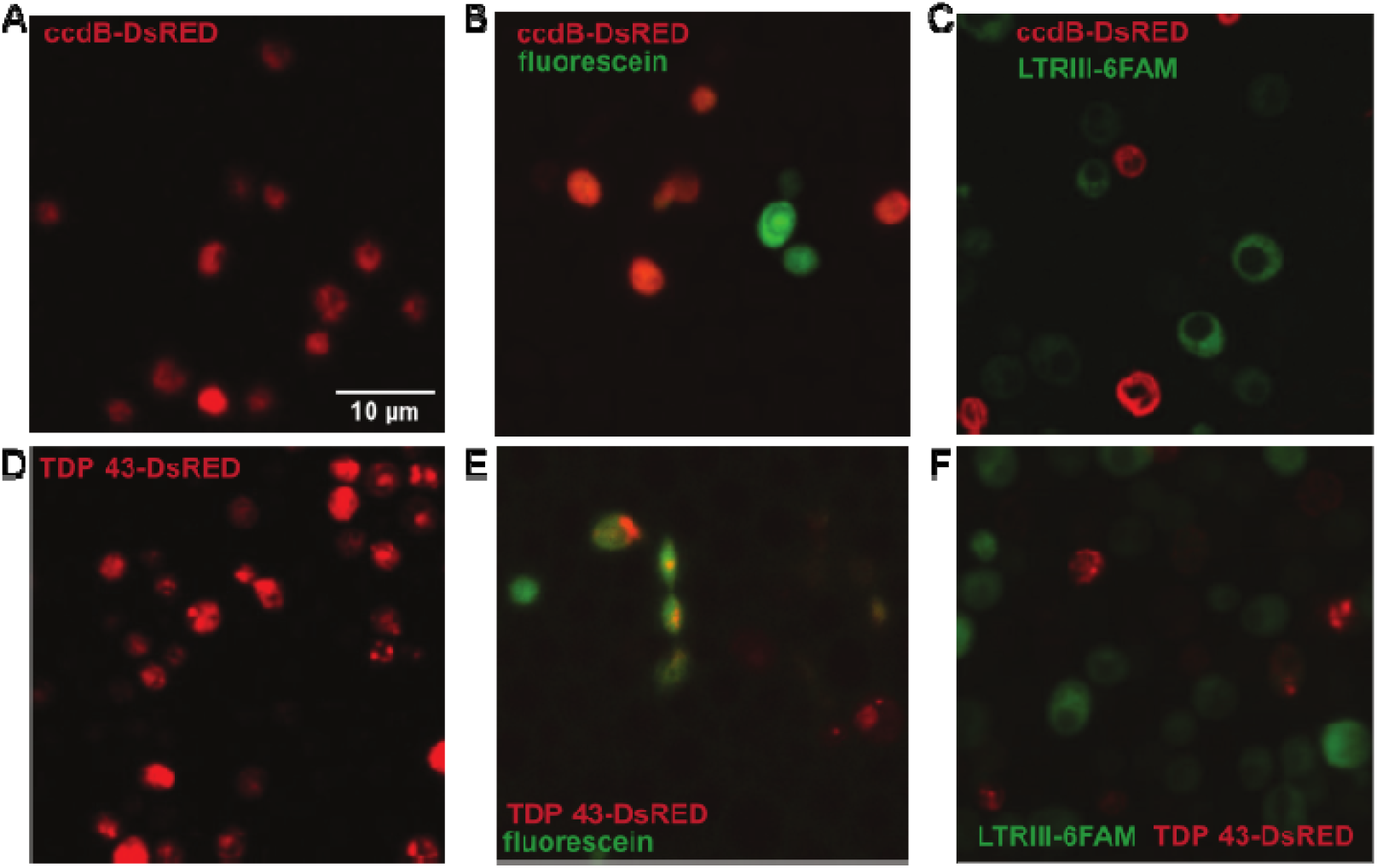
G-quadruplexes and fluorescein (green) enter the yeast cytoplasm when added to the yeast media. A. ccdB-DsRED expressing yeast, with no added rG4s or fluorophore dyes confocal microscopy. B. ccdB-DsRED yeast with 10 µM fluorescein. c. ccdB-DsRED yeast with 10 µM LTRIII-6FAM. D. TDP43-DsRED expressing yeast, with no added rG4s or fluorophore dyes confocal microscopy. E. TDP43-DsRED yeast with 10 µM fluorescein. F. TDP43-DsRED yeast with 10 µM LTRIII-6FAM.

**Supplementary Figure 6.**
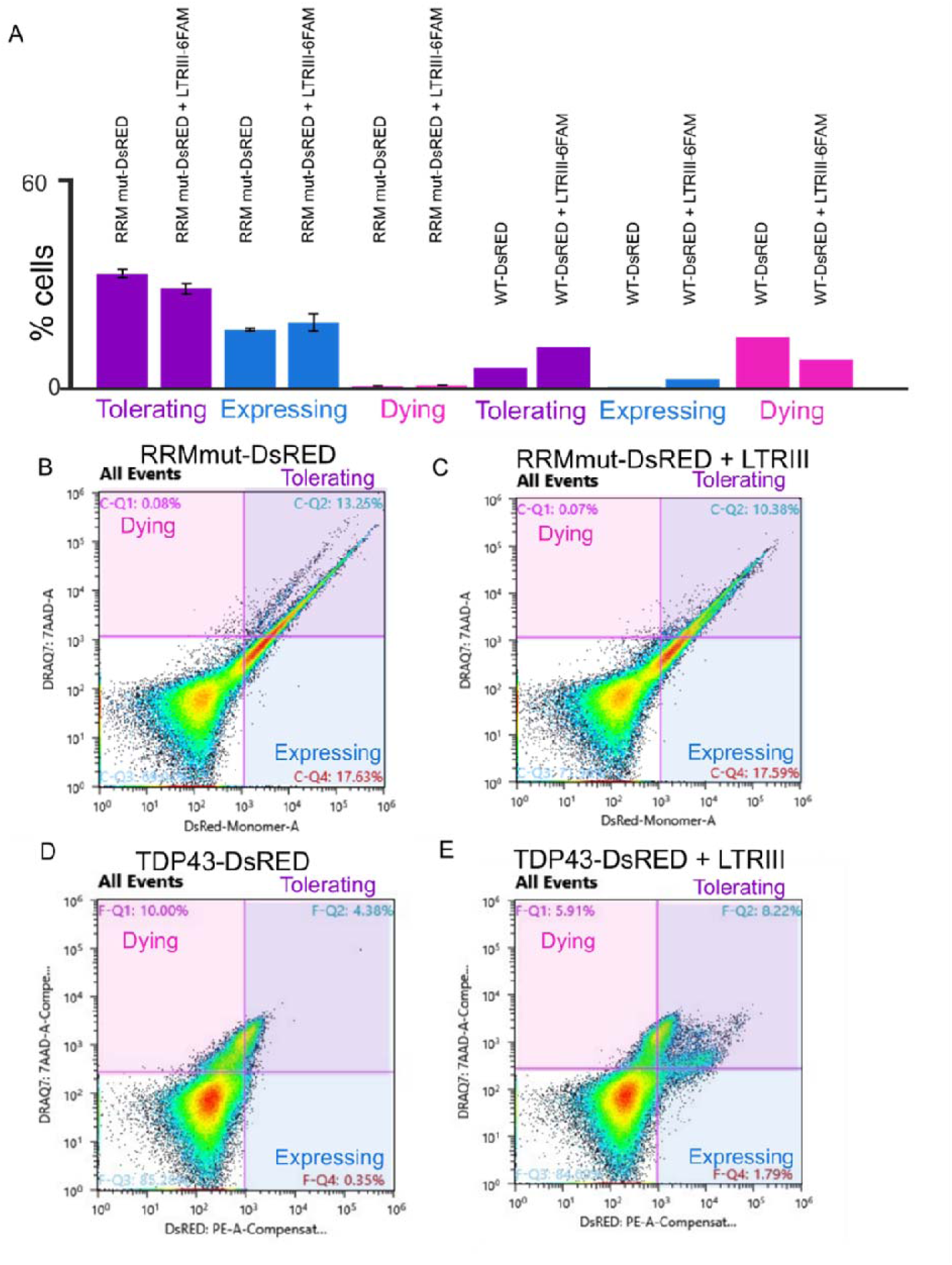
5F->L RRM mut TDP43 is not cytotoxic in yeast, in the presence and absence of G4 LTRIII-6FAM. A. Quantifications of the tolerating, expressing, and dying populations of yeast cultures expressing WT TDP43-DsRED and RRM mut (5F->L)-DsRED in the presence and absence of LTRIII-6FAM. B-E. Representative flow cytometry of the B. 5F->L TDP43 RRM mut - DsRED C. 5F->L TDP43 RRM mut -DsRED with LTRIII-6FAM D. WT TDP43-DsRED and E. WT TDP43-DsRED with LTRIII-6FAM stained with DRAQ7 showed little change between the presence and absence of G4 LTRIII-6FAM, and was not cytotoxic; n = 3.

**Supplemental Figure 7.**
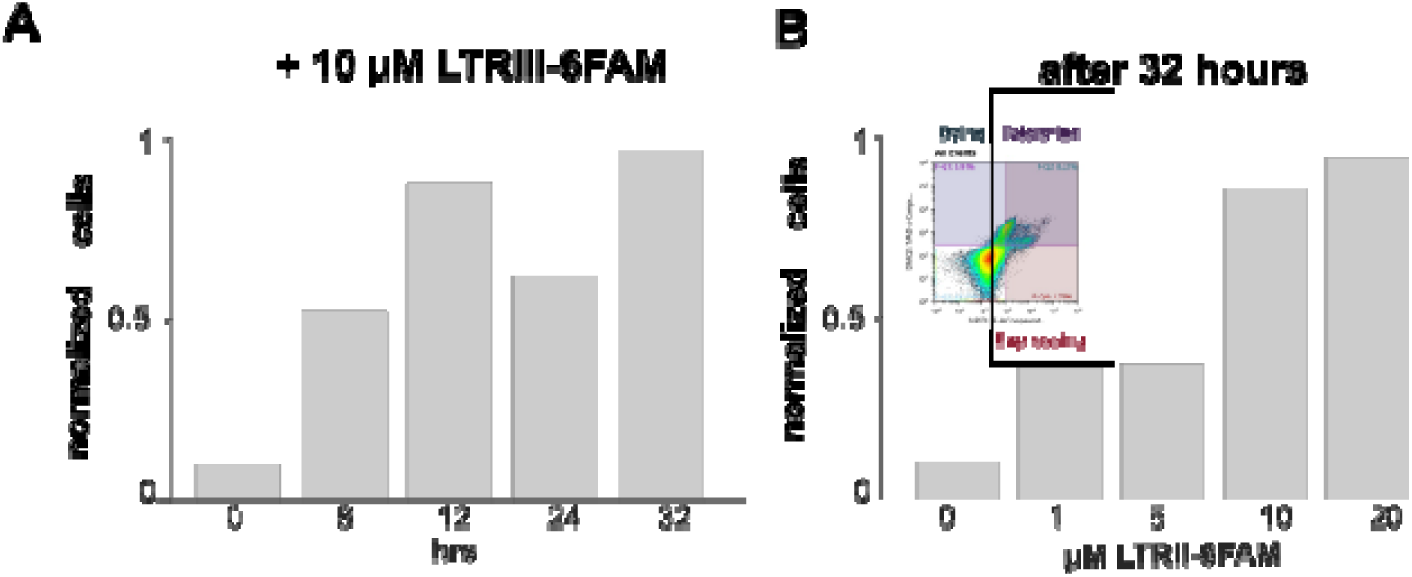
A replicate of Figure 1 C, D. A. A time course of the normalized % cell tolerating and expressing TDP43-DsRED per 100,000 yeast cells based on DsRED and DRAQ7 intensity. B. A dosage curve of the normalized % cells tolerating and expressing TDP43-DsRED per 100,000 yeast cells based on DsRED and DRAQ7 intensity.

**Supplement Figure 8.**
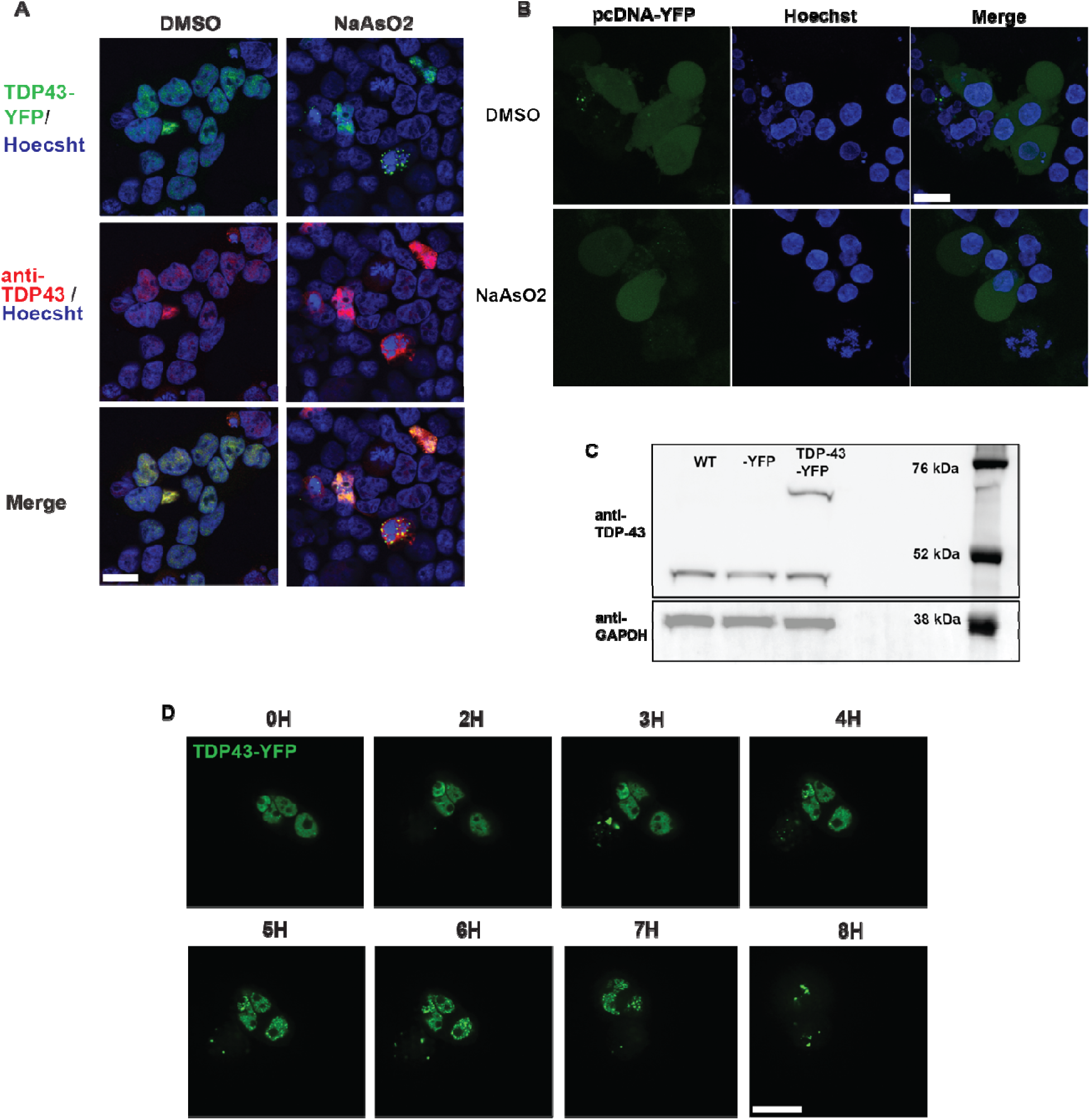
Oxidative stress induction in HEK293T cells with stable TDP43 overexpression. A. Immunostaining shows colocalization of stable overexpressing HEK-TDP43-YFP (green) and anti-TDP43, Alexa-fluor 647 (red), Hoescht nuclear label (blue), 4% PFA, 40x, scale bar = 20 µM. B. Live cell imaging of empty vector pcDNA-YFP was transfected into HEK cells and addition of NaAsO_2_ showed no condensation of off target fluorescence. C. Western blot analysis of 10 µg HEK cell lysates (WT, -YFP, -TDP43-YFP), stained for anti-TDP43 and GAPDH. D. Dose-dependent live confocal imaging was taken over 8 hrs. When treated with 50 µM of NaAsO_2_, nuclear and cytoplasmic TDP43 inclusions are induced in HEK-TDP43-YFP model in as little as 4 hrs. To lessen cell death triggered by NaAsO_2_, concentration was dropped to 15 µM.

**Supplementary Figure 9.**
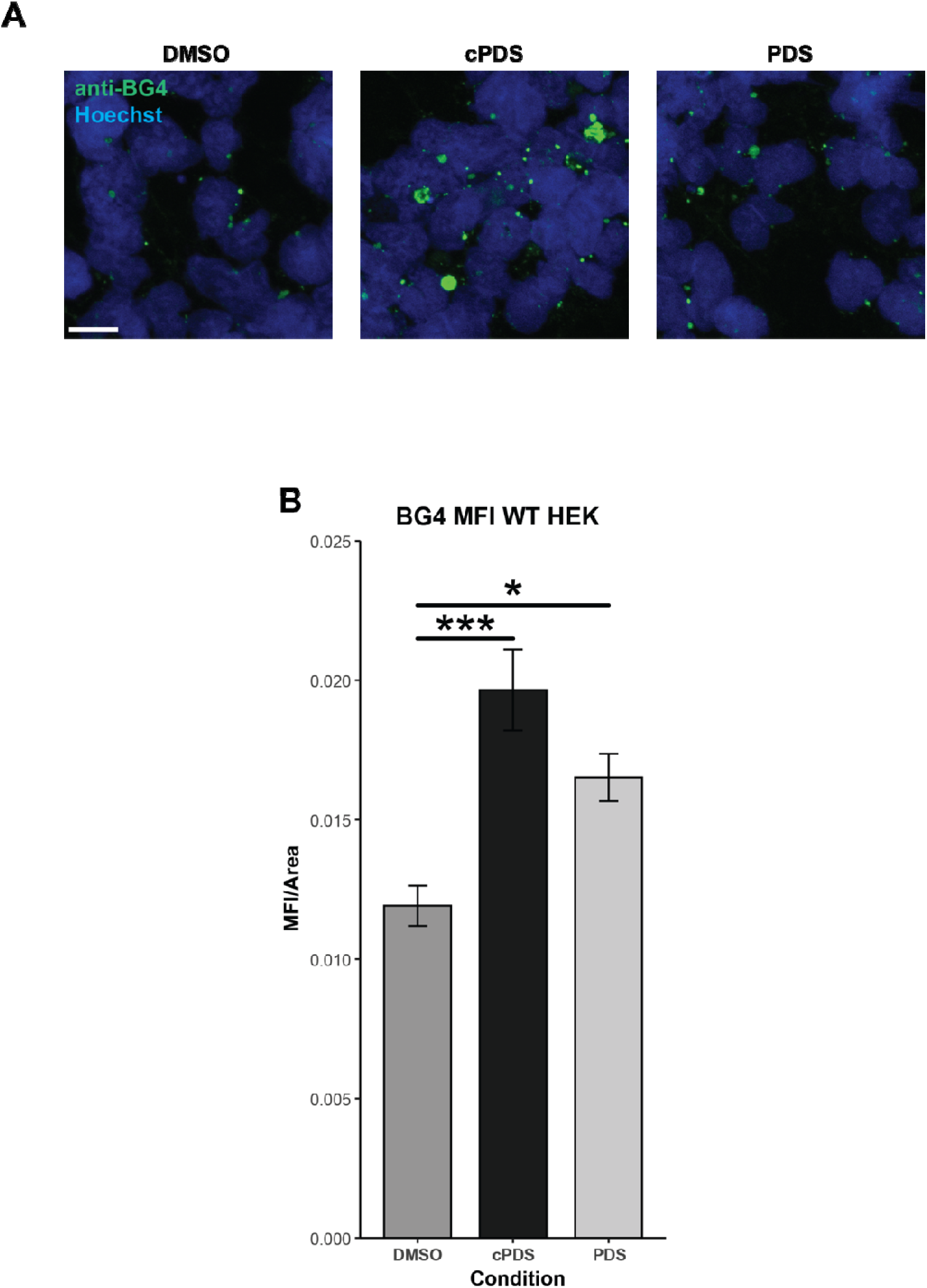
Monitoring the effects of G4 binders on G4 levels in WT HEK cells. A. Immunofluorescence images of HEK293T WT cells treated with the G4 binders cPDS and PDS, or the DMSO vehicle control. Cells were fixed in 4% PFA and stained with anti-BG4 (AF488) to visualize G4 structures, and nuclei were labeled with Hoechst. Treatment with cPDS resulted in a greater accumulation of G4s compared to PDS or the DMSO control, scale bar = 20 µM. B. Quantification of mean fluorescence intensity (MFI) of anti-BG4 staining under different treatment conditions. Statistical significance: ***, *p* < 0.001; *, *p* = 0.027.

**Supplemental Figure 10.**
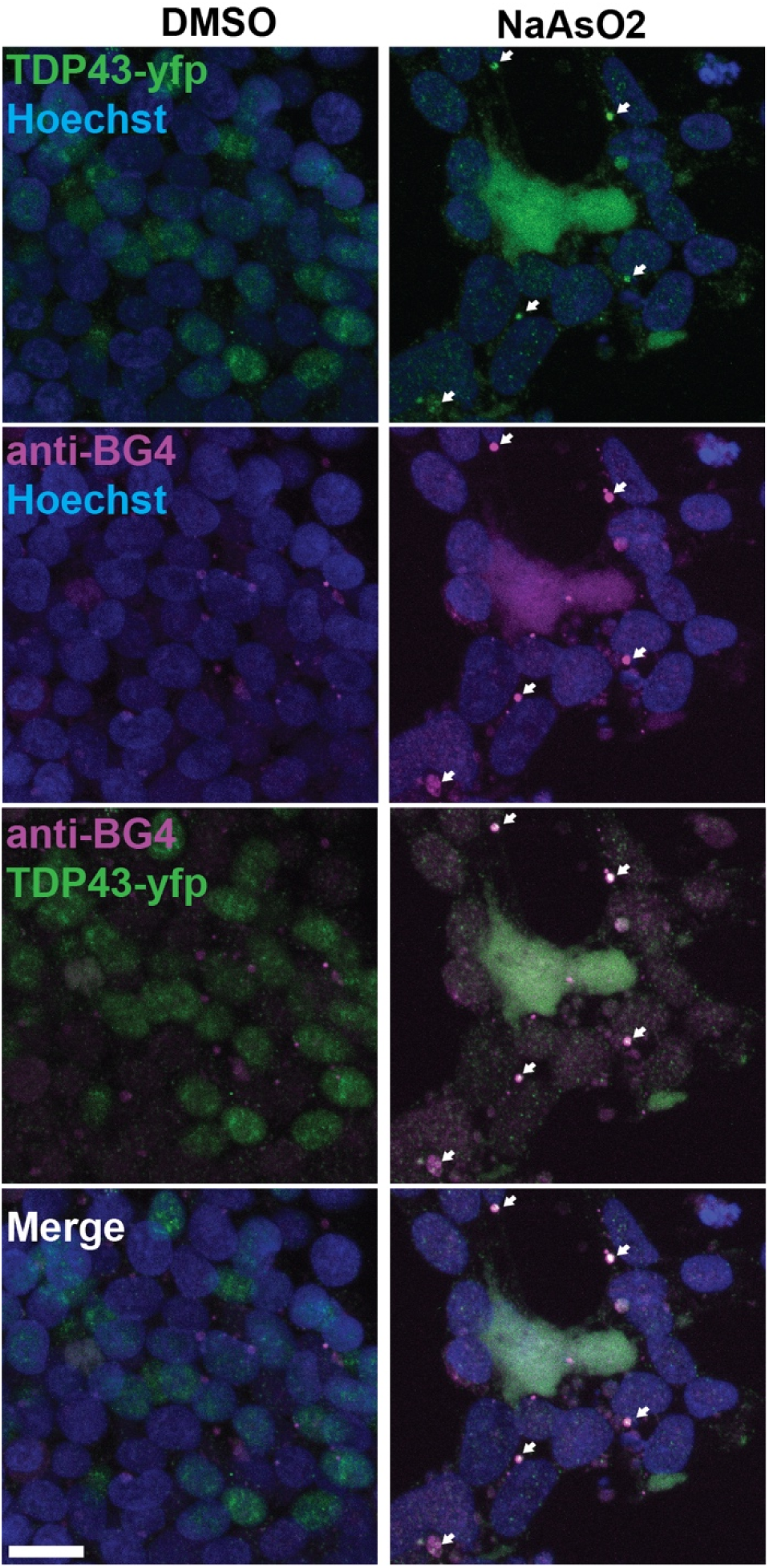
BG4 colocalizes to TDP43 condensates in HEK cells under oxidative stress. HEK-TDP43-YFP exposed to NaAsO_2_ fixed in 4% PFA and stained for BG4 (AF647) show cytoplasmic rG4 overlap with TDP43 condensates. Scale bar = 20 µM

**Supplemental Figure 11.**
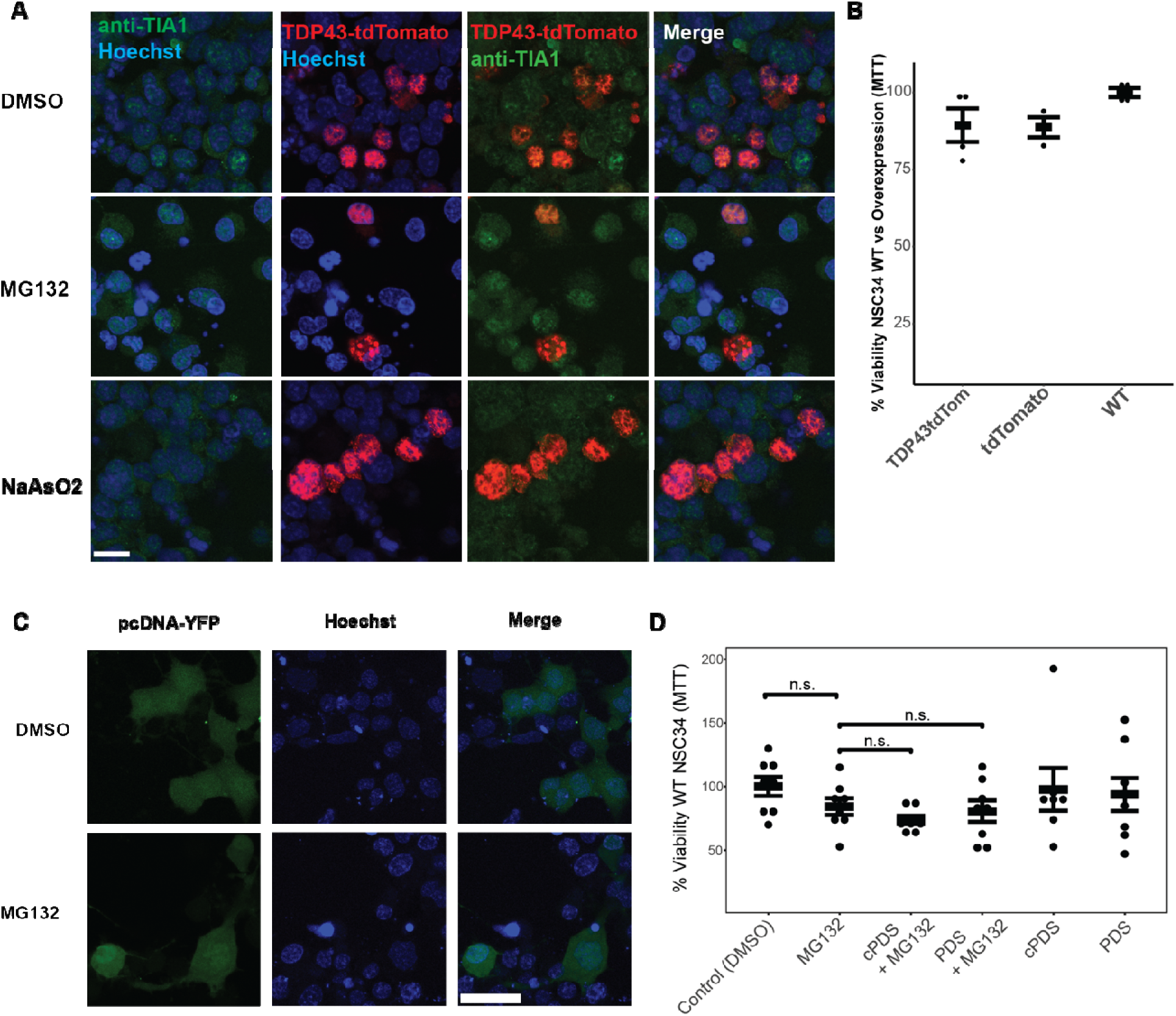
WT NSC34 and transfected pcDNA -YFP show insignificant cell death or condensation under proteasomal stress. A. ICC of NSC34-TDP43-tdTomato cells stressed, fixed in 4% PFA, and stained against stress granule marker, TIA-1. B. NSC34 MTT comparing cell viability among WT, tdTomato, and TDP43-tdTomato overexpression, n = 4. C. Live cell imaging of overexpression empty vector pcDNA-YFP in NSC34 cells unremarkable after addition of proteasome inhibitor, scale bar = 40 µM. D. WT NSC34 MTT in the presence of G4 binders PDS and cPDS and MG132, n = 6.

**Supplementary Figure 12:**
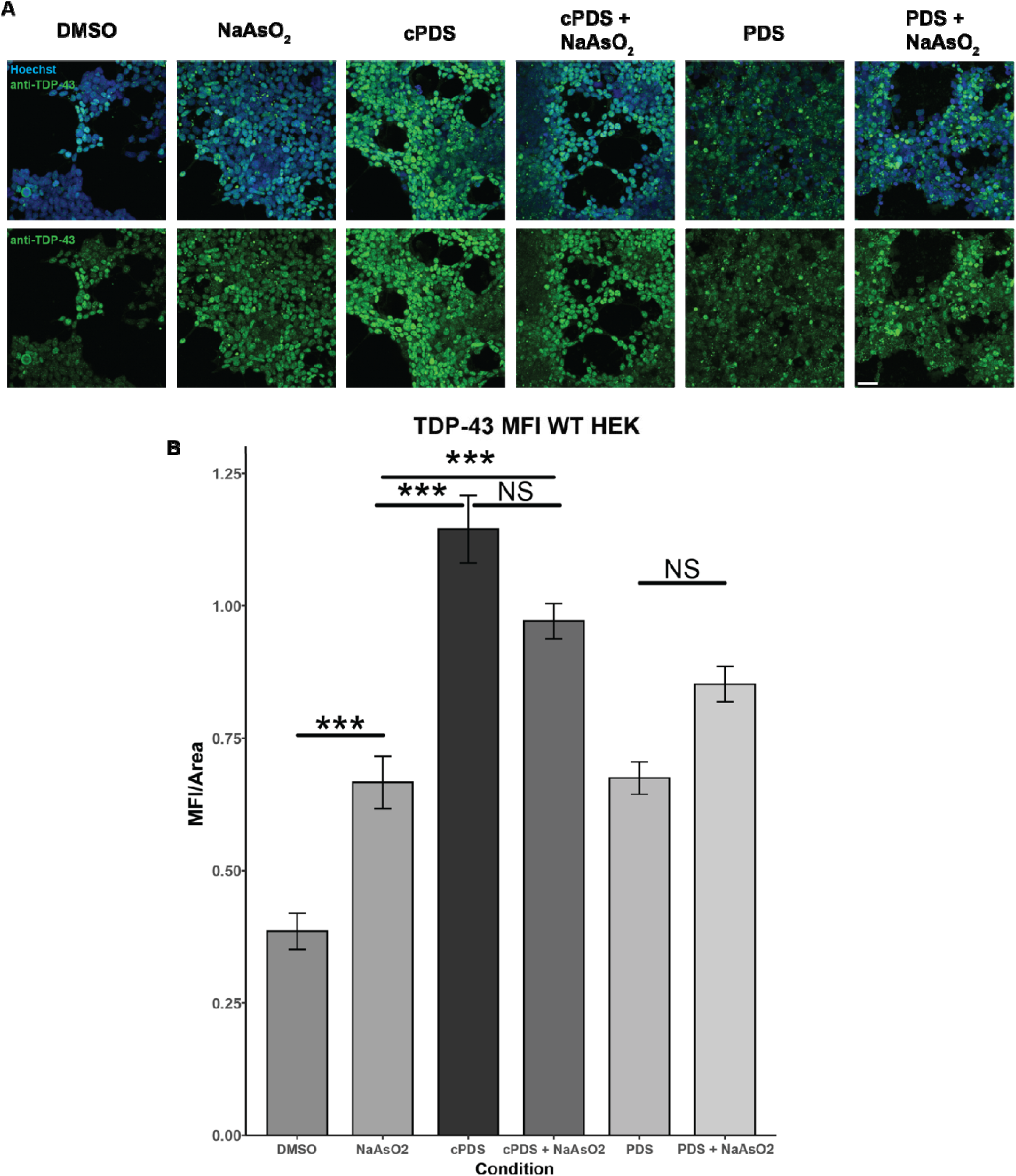
Endogenous TDP43 signal increases upon exposure to G4 binders prior to stress. A. ICC of WT HEK293T cells were fixed in 4% PFA and stained with anti-TDP43 to detect endogenous TDP43 structures, showing increased TDP43 levels in the presence of G4 binders, scale bar = 40 µM. B. Quantification of mean fluorescence intensity confirms this increase. ***, p < 0.001.

## Notes

### Competing Interest Statement

The authors have declared no competing interest.

### Summary of Updates

New experiments and controls for several figures.

